# Common origin for effector and regulatory follicular and tissue-Adapted CD4+ T cells in Non-Small Cell Lung Cancer

**DOI:** 10.1101/2025.09.17.674622

**Authors:** Jimena Tosello Boari, Wilfrid Richer, Yoann Missolo-Koussou, Elisa Bonnin, Sebastien Lemoine, Marco Antônio Pretti, Rodrigo Nalio Ramos, Jordan Denizeau, Sylvain Baulande, Mylène Bohec, Sonia Lameiras, Philippe Martin, Rafael Mena Osuna, Jeremy Mesple, Wenjie Sun, Marine Lefevre, Agathe Seguin-Givelet, Christine Sedlik, Filipa Ramos, Saumya Kumar, Leïla Perié, Luis Graça, Olivier Lantz, Edith Borcoman, Nicolas Girard, Joshua J Waterfall, Eliane Piaggio

## Abstract

Tumor-invaded lymph nodes (LNs) serve as critical hubs for anti-tumor immunity, yet their role in orchestrating immune responses remains poorly understood. Using integrated single-cell RNA sequencing, T cell receptor sequencing, and chromatin accessibility profiling, we analyzed CD4+ T cells from matched blood, tumor-invaded LNs, and tumors of treatment-naïve non-small cell lung cancer patients. We identified distinct immunological landscapes across these compartments. Compared to blood, tumor-invaded LNs and tumors were enriched for follicular regulatory T cells (Treg-Tfr), conventional T cell subsets with Tfh-like characteristics (Tconv-Tfh and Tconv-CXCL13), and tissue-resident memory Tregs (Treg-Trm). These populations share a BATF-dependent transcriptional program that governs T-cell activation and tissue adaptation, while simultaneously engaging distinct, subset-specific regulatory networks. Integrative TCR-RNA analysis revealed that tumor-reactive, neoantigen-specific T cell clones were enriched within these subsets and demonstrated extensive LN-tumor clonal sharing, indicating active recirculation between compartments. Through clonal coupling analysis and trajectory inference, we uncovered that Treg-Tfr cells function as multipotent progenitors that bifurcate into tissue-resident Treg-Trm or into ex-Tregs adopting a Tfh-like CXCL13+ effector phenotype. Remarkably, follicular CD4+ T subsets from LNs and tumor were transcriptionally and epigenetically similar and localized to analogous germinal center niches. These findings establish tumor-invaded LNs as functional extensions of the tumor microenvironment that generate and maintain tumor-reactive CD4+ lineages. The identification of tissue-resident Treg-Tfr plasticity reveals a critical developmental checkpoint that could be therapeutically targeted to redirect immunosuppressive programs toward anti-tumor effector responses.

## INTRODUCTION

CD4+ T cells orchestrate adaptive immunity through functional plasticity, differentiating into specialized subsets that either promote or suppress anti-tumor responses.^1^ This cellular diversity emerges within interconnected lymph nodes (LNs) and tumors, both serving as sites for T cell priming and differentiation.^2–5^ The dynamic trafficking of CD4+ T cells between these compartments creates an immunological circuit fundamental to cancer surveillance and response.^6–10^ However, this anatomical network also facilitates metastatic spread, with LNs frequently representing the first sites of tumor colonization.^2,11–13^ Invaded LNs constitute unique immunological environments where tumor cells and immune cells coexist, potentially creating specialized niches that shape CD4+ T cell fate decisions differently than either blood or primary tumors. The balance between effector and regulatory immunity within invaded LNs and their relationship to tumor immunity remains poorly understood.

Within the tumor microenvironment, CD4+ T cells show high functional diversity.^14–16^ Recent advances in single-cell technologies and animal models have extended this heterogeneity beyond traditional T helper classifications. Among conventional CD4+ T cell (Tconv) populations, follicular T cells are key regulators of anti-tumor immunity through their participation in GC responses.^17,18^ T follicular helper (Tfh) cells, characterized by PD-1+ICOS+CXCR5+ expression, BCL6-driven transcriptional programs, and IL-21 production, represent the prototypical follicular effectors that initiate GC responses.^19^ Additionally, the follicular T cells include other specialized populations, such as CXCL13-producing T cells mediating tumor-specific responses, ^20–25^ tissue-resident helpers supporting local B cell maturation,^26^ and peripheral helpers driving TLS formation.^27^ Tumor-infiltrating Tregs exhibit pronounced tissue adaptation, functioning as tissue-resident Tregs (Treg-Trm) to counterbalance effector responses^28–33^. The regulatory repertoire also includes FOXP3+ T follicular regulatory (Tfr) cells controlling GC reactions, ^34–39^ and other potential actors such as IL-10-producing Tfr variants,^40^ Tr1 cells,^41^ or FOXP3+ cells emerging during GC contraction,^40^ among others.^42,18,24,37,43^ Intriguingly, these regulatory populations often display Tfh-like features^44,31^, potentially enabling precise spatial positioning for targeted immunosuppression. Despite extensive characterization of tumor-infiltrating CD4⁺ T cells, critical questions remain. What is the overall landscape of CD4⁺ T cell populations in invaded LNs compared with those in tumors and blood? What are the developmental relationships among the full spectrum of follicular subsets? How are CD4⁺ T cells in invaded LNs related to their counterparts in tumors, and what roles do they play in shaping anti-tumor immunity?

To address these questions, we performed comprehensive multi-omic profiling of CD4+ T cells across matched blood, invaded LNs, and tumors from treatment-naïve NSCLC patients. Our integrated single-cell multi-omics analyses revealed that invaded LNs harbor distinct T cell subpopulations that mirror intra-tumoral CD4+ diversity rather than peripheral blood. We identified specialized follicular regulatory populations and tissue-adapted subsets sharing core transcriptional programs of T cell activation and tissue residency across LNs and tumors, with tumor-reactive clones actively trafficking between these compartments. Unexpectedly, we discovered developmental plasticity within the regulatory compartment, where Treg-Tfr cells can bifurcate into tumor resident Treg or Tfh-like CXCL13+ Tconvs under local microenvironmental cues. These findings redefine invaded LNs as active immunological partners with tumors, maintaining shared CD4+ T cell populations and providing niches for both regulatory and effector differentiation. Our work reveals how the invaded LN-tumor axis shapes anti-cancer immunity and identifies regulatory T cell plasticity as a potential target to enhance immunotherapy.

## RESULTS

### Regulatory and Conventional Follicular CD4+ T Cell Subsets Accumulate in Invaded Lymph Nodes and Tumors

Although several follicular and regulatory CD4⁺ T cell populations have been identified within tumors, the landscape of CD4⁺ T cells residing in invaded LNs, remains poorly defined. To address this, we performed scRNA-seq coupled to TCR-seq on CD4+ T cells isolated from matched reactive LNs (suspected of being invaded at the time of the surgery), tumor, and blood from treatment-naïve patients. Tregs and Tconvs from each location were FACS-sorted as CD25^high^CD127^low^ and CD25^low^CD127^high/middle^ CD45+CD4+ live cells, respectively. As Tregs are typically less abundant than Tconvs, we mixed the two populations at a 1:1 Treg:Tconv ratio prior to loading the 10x Genomics RNA/TCR Immunoprofiling kit, thereby increasing the statistical power for Treg subset characterization (**Fig. 1A**).

**Fig. 1.**
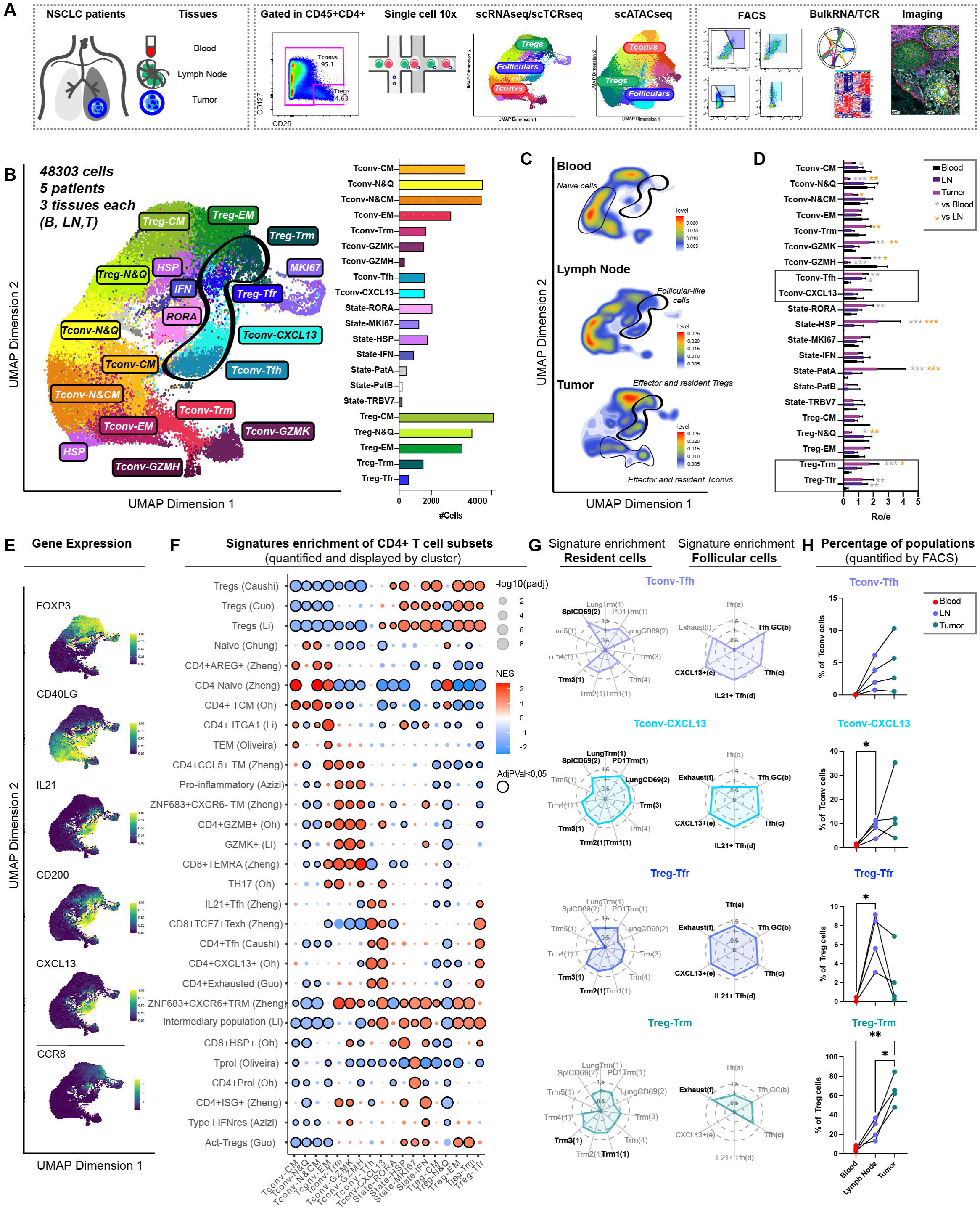
scRNA-seq landscape of paired blood, lymph node (LN), and tumor samples from NSCLC patients. **(A)** Schematic of the study design. CD4⁺ Tregs and Tconvs were FACS-purified from matched blood, LN, and tumor samples from NSCLC patients, then combined in equal proportions per tissue and loaded onto 10X single-cell chips for scRNA-seq, TCR-seq, and scATAC-seq analysis. Additional analyses included FACS, imaging, bulk RNA/TCR sequencing, and functional assays. **(B)** UMAP projection of cells from the integrated scRNA-seq dataset (three tissues, five patients, 15 samples). Each dot represents an individual cell, colored by cluster identity and classified into three groups: Tregs, Tconvs, and Treg/Tconv state-driven clusters. The accompanying bar plot shows the number of cells per cluster. **(C)** Umap plots representing the cell density gradients by tissue. **(D)** Bar plot showing cluster size distribution across tissues, represented as the ratio of observed cell numbers to random expectation (RO/E), calculated using a chi-square test (*p < 0.05; **p < 0.01; ***p < 0.001; paired Student’s t-test). **(E)** UMAP plots displaying normalized expression levels of Treg-, Tconv-, follicular-, and Treg resident memory related genes (blue: low expression; yellow: high expression). **(F)** GSEA enrichment analysis. Dot plots showing the Normalized Enrichment Score (NES) (color intensity) and adjusted p-value (circle size) for each signature. Signatures with an adjusted p-value < 0.05 are outlined in black, while non-significant ones are unmarked. **(G)** Spider plots depicting the Normalized Enrichment Score (NES) of follicular (upper plots) and tissue-resident (lower plots) signatures derived from publicly available datasets. Signatures with p < 0.05 are bolded, while non-significant ones are shown in gray. Signature references: (a) Massoni-Badosa 2024: Tfr; (b) Massoni-Badosa 2024: Tfh-GC; (c) Caushi et al. 2021: Tfh; (d) Zheng et al. 2023: IL21⁺Tfh; (e) Oh et al. 2020: CXCL13⁺; (f) Guo et al. 2018: Exhausted; (1) Clarke et al. 2019: PD1Trm, Trm1-5, Lung Trm; (2) Kumar et al. 2017: SplCD69 and LungCD69; (3) Wu et al. 2020: Trm; (4) Cheuk et al. 2017: Trm. **(H)** Scatter plots quantifying the percentage of indicated populations within total Tregs or Tconvs across tissues. FACS analysis: n = 4 LNs from NSCLC patients. Statistical analysis: paired one-way ANOVA with Tukey’s multiple comparisons post hoc test (* = p ≤ 0.0332; ** = p ≤ 0.0021; *** = p ≤ 0.0002; **** = p ≤ 0.0001).

For the single-cell transcriptomic analysis we studied CD4+ T cells from five individuals (three tissues in each patient, for a total of 15 samples (**Fig. 1A** and **Table1**). The transcriptome of 48,303 cells was recovered in total after quality control. Cells from all patients and tissues were integrated in a single dataset (**Fig. S1A-C**). Using unsupervised graph-based clustering we identified 21 clusters visualized in the Uniform Manifold Approximation and Projection (UMAP) plot, along with the corresponding cell counts per cluster in **Fig. 1B**. Although the relative proportions of these subpopulations varied across tissues, the majority of clusters were represented in all tissues and patients (**Fig. S1B-C)**.

Based on the analysis of clustering resolution hierarchy (**Fig. S1D**) and characteristic gene and signature expression (**Fig. 1E-F, S1E-G** and **Table 2-3**), cell clusters were classified as Tregs, Tconvs, or clusters containing both Tregs and Tconvs grouped by their functional state rather than lineage identity. The five Treg clusters expressed high levels of *FOXP3* and *IL2RA* transcripts and were enriched in Treg signatures. The nine Tconv clusters expressed high levels of *CD40LG* and *IL7R* transcripts. The four “State” clusters presented a mix of Treg and Tconv cells (**Fig. 1E-F** and **S1E-G**).

In more detail, the five Treg clusters segregated as: naïve & quiescent (**Treg-N&Q**; expressing *TCF7* and *CCR7*), central memory (**Treg-CM**; *SELL and S100A4*), effector memory (**Treg-EM**; *HLA-DR and CXCR3*), effector and tissue-resident (**Treg-Trm**; *CCR8, TNFRSF9, and IL1R2*), and follicular Tregs (**Treg-Tfr**; *CXCL13, ICA1,* and *IL1R2*). The nine Tconv clusters were identified as: naïve & quiescent (**Tconv-N&Q**; *TCF7*, *SELL and RPS6*), naïve & central memory (**Tconv-N&CM**; *S1PR1, ANXA1,* and *GPR183*), central memory (**Tconv-CM**; *S100A4*, *AREG, TCF7*), effector memory (**Tconv-EM**; *ANXA1, and NFKBIA*), the effector populations TEMRA/innate-like (**Tconv-GZMH;** *GZMH,* and *GNLY*) and terminal-effector (**Tconv-GZMK;** *GZMK,* and *EOMES*), one tissue-resident (**Tconv-Trm;** (*GZMB* and *ITGAE),* and two follicular helper clusters: T follicular helper (**Tconv-Tfh**; *ICOS*, *CXCR5, and BCL6*) and an inflammatory Tfh subset (**Tconv-CXCL13**; *CXCL13*, *KLRB1,* and *IL21).* The main state clusters were identified as: cycling (**State-MKI67**; *MKI67*, *TOP2A*), activated (**State-RORA**; *RORA, STAT3*), IFN response (**State-IFN**; *IFIT1, ISG15*), and stress response (**State-HSP**; *HSPE1, DNAJB1*).

Analysis of the tissue origin of the cells revealed distinct cluster distribution patterns across tissues (**Fig. 1C**). Notably, effector and terminally differentiated subsets were enriched in tumors and absent in LNs and blood, while follicular subsets accumulated in both LNs and tumors compared to blood.

More specifically, quantification of cells per cluster across the three tissues, based on observed versus expected proportions (**Fig. 1D**), revealed that N&Q and CM cells were preferentially found in blood and LNs, whereas the state clusters HSP and RORA, along with terminally differentiated effector Tconv-GZMK cells, accumulated primarily in tumors. Both Treg-Tfr and Tconv-Tfh follicular clusters were significantly enriched in LNs and tumors, while tissue-resident populations (Treg-Trm and Tconv-Trm) were predominantly localized to tumors. Consistent with this distribution, follicular CD4⁺ subsets and Treg-Trm displayed gene expression signatures characteristic of both follicular and tissue-resident programs (**Fig. 1G** and **S1H**). Within this spectrum, the Tconv-CXCL13 cluster most strongly combined residency and follicular features, despite its heterogeneous distribution across tissues. As expected, classical residency signatures were highly enriched in Tconv-Trm, whereas naïve and quiescent Tregs and Tconvs lacked these programs entirely.

To validate these results in more patients we performed FACS analysis. We identified novel, differentially expressed surface markers characterizing these subpopulations, with a particular focus on Tfh-like subsets and Treg-Trm. As shown in **Fig. S1F-G**, CD200, PDCD1, and BTLA collectively distinguish follicular cells from other Tconv and Treg populations; IL2RA (CD25) enables Treg identification; CCR8 and ICOS differentiate Treg-Trm cells; high CD200 and BTLA expression identified Treg-Tfr cells; and CXCR5 and KLRB1 are differentially expressed between Tconv-Tfh and Tconv-CXCL13 subsets. Based on these markers, we designed a FACS sorting strategy incorporating CD45RA to exclude naïve cells and CD127 to better distinguish Tregs from Tconvs (**Fig. S1I**). We further validated the identity of FACS-sorted populations through bulk RNA sequencing of follicular subsets and Treg-Trm, using naïve cells as controls. Cells were isolated from LNs and tumors of five additional NSCLC patients. Clustering of the bulk RNA-seq data reflected cell type rather than tissue origin (**Fig. S1J, right**), and these populations were significantly enriched for genes and signatures derived from the single-cell RNA-seq analysis of the corresponding clusters (**Fig. S1J, left; Table 4**).

Using this strategy, we quantified the distribution of CD4⁺ subpopulations across blood, LNs, and tumors from additional NSCLC patients (**Fig. 1H**). Consistent with scRNA-seq results, follicular subsets were enriched in LNs, particularly Tconv-Tfh and Tconv-CXCL13, whereas Treg-Tfh and Treg-Trm cells showed gradual accumulation in both LNs and tumors.

Overall, integration of scRNA-seq and cytometry analyses indicates that invaded LNs contain a diverse repertoire of CD4⁺ T cells with distinct transcriptomic features and tissue-specific distributions. Follicular CD4⁺ T cells and Treg-Trm populations are present in both LNs and tumors, suggesting that these subsets may regionally shape tumor-associated immune responses.

### Follicular CD4⁺ T Cells Share Gene Regulatory Programs of TCR Activation and Tissue Residency with Trm Cells

Follicular and tissue-resident CD4⁺ T cell subsets display distinct transcriptomic profiles; however, their epigenetic programs and the specific regulatory patterns that define their functional states remain incompletely characterized. To address this gap, we performed scATAC-seq on CD4+ T cells from two additional NSCLC patients, using the same sorting strategy as for scRNA/TCRseq (**Fig. 1A**).

We analyzed the chromatin accessibility landscape of a total of 50,171 cells passing the quality controls (**Fig. S2A-C**). All scATAC-seq samples were integrated into a single dataset and subjected to unsupervised graph-based clustering, which identified 17 accessibility subsets (**Fig. 2A, S2D, Tables 5-6).** Of note, the number of differentially accessible genes within scATAC-seq clusters, both overall and per cluster, was four-times higher than those detected in the scRNA-seq analysis. **Fig. 2B** and **S2C** show the distribution of nuclei across clusters, tissues, and patients.

**Fig. 2.**
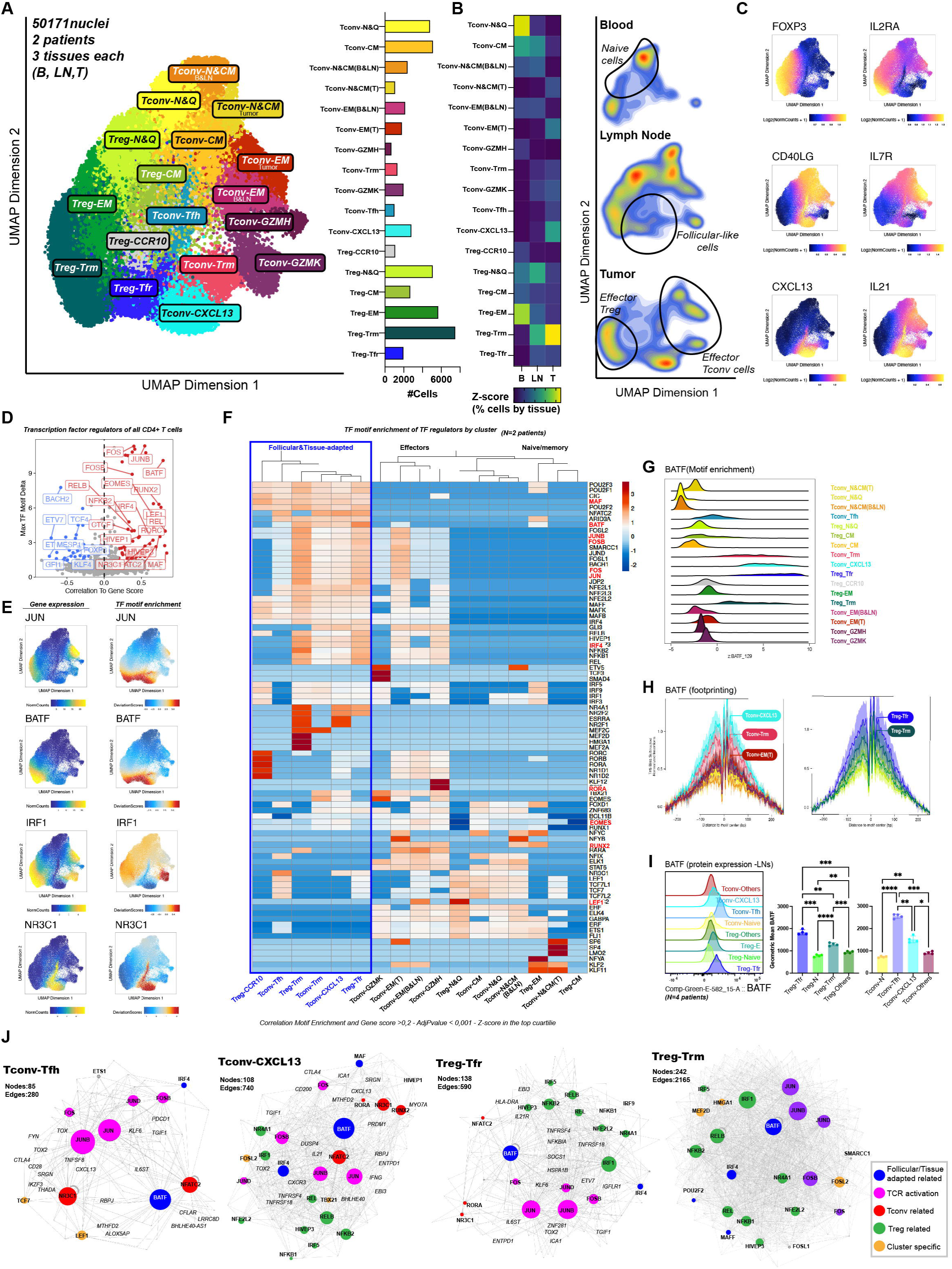
Single-cell ATAC landscape of paired blood, LN, and tumor samples from NSCLC patients. **(A)** UMAP projection of integrated scATAC-seq nuclei from six samples (three tissues, two patients). Dots represent individual nuclei, colored by cluster identity, classified into two groups: **Tregs** and **Tconvs**. **Right:** Bar plot indicating the number of nuclei per cluster. **(B) Left:** Heatmap displaying the relative proportion of each cluster per tissue: blood (B), lymph node (LN), and tumor (T). **Right:** Density gradients showing per-tissue distribution of cells in UMAP space. **(C)** UMAP plots displaying gene score levels (blue = low, yellow = high) for six key genes defining Tregs, Tconvs, and follicular cells. **(D)** Volcano plot illustrating maximum TF motif delta (y-axis; TF motif deviation score driving cluster variation, motif delta ≥ 2) vs. the correlation value between TF motif enrichment and gene expression (x-axis; r ≥ 0.2 for positive regulators). The top 25% most variable TFs are color-coded: red (positive regulators), blue (negative regulators). **(E)** UMAP projections showing **left:** gene expression (scRNA-seq and scATAC-seq integration) and **right:** ChromVAR TF deviation scores (TF motif enrichment) for selected TFs. **(F)** Heatmap of TF motif enrichment (FDR < 0.05, ME > 1) for inferred positive regulators from (D) across clusters, hierarchically clustered. Key selected TFs are highlighted in red. **(G-H)** Visualization of BATF motif enrichment deviation scores **(G)** and footprint analysis **(H)** across clusters. **(I)** FACS analysis. **Left:** Histograms of BATF expression in gated populations from LNs. **Right:** Bar plots quantifying BATF geometric mean fluorescence intensity in subsets from a representative NSCLC LN sample (n = 4 patients). Statistical test: One-way ANOVA (paired) with Tukey’s multiple comparisons post hoc test (* = p ≤ 0.0332; ** = p ≤ 0.0021; *** = p ≤ 0.0002; **** = p ≤ 0.0001). **(J)** TF-target gene network for the three follicular clusters and Treg-Trm cells, showing positive regulators from (D). Node size correlates with out-degree (number of target genes), and node color represents association with TCR activation, tissue-imprinted programs, Tconv/Treg identity, or cluster specificity (see legend).

Similar to scRNA-seq datasets, the 17 subsets were classified as Tregs or Tconvs based on the accessibility of the *FOXP3/IL2RA* and *CD40LG/IL7*R loci, respectively (**Fig. 2C**). Moreover, follicular cells, including Treg-Tfr, Tconv-Tfh and Tconv-CXCL13, were identified by the accessibility of the *CXCL13* and *IL21* loci (**Fig. 2C**).

More specifically, Treg clusters included: naïve & quiescent (**Treg-N&Q**: marked by *LEF1* and *CCR7*), central memory (**Treg-CM**: *SELL* and *AREG*), effector memory (**Treg-EM**: *HLA-DR* and *TNFRSF4*), effector and tissue resident (**Treg-Trm**: *TNFRSF9*, *CCR8, CD80*), and a minor ***CCR10*** group. Tconv clusters comprised: naïve & quiescent (**Tconv-N&Q**: characterized by *SELP*, *NOSIP* and *SELL*); central memory (**Tconv-CM**: *AREG,* and *ANXA1*); two naïve & central memory clusters, - one enriched in blood and LNs and another in tumors (**Tconv-N&CM(B&LN)** and **Tconv-N&CM(T)**: *S1PR1,* and *TCF7*); two effector memory clusters (**Tconv-EM(B&LN)** and **Tconv-EM(T)**: *ANXA1, NFKBIA,* and *TNF*), and three effector Tconv clusters (tissue-resident **Tconv-Trm:** *GZMB, CCL3,* and *ITGAE;* TEMRA/innate like **Tconv-GZMH**: *GZMH,* and *GNLY*; and terminal-effector (**Tconv-GZMK**: *GZMK, LAG-3,* and *EOMES*). Follicular clusters included follicular Tregs (**Treg-Tfr;** *TCF7, IL1R2, TOX*), follicular Tconvs (**Tconv-Tfh**: *ICOS, CXCR5,* and *BCL6*), and inflammatory follicular Tconvs (**Tconv-CXCL13**; *CXCL13, IL21,* and *IL17A*). A detailed characterization of each cluster is provided in **Fig. S2E-F** and **Table 5).**

Overall, we found similar subpopulations to those in the scRNA-seq dataset, except for the state-driven clusters which were absent in the scATAC-seq characterization. This was further confirmed by integrating scATAC-seq and scRNA-seq datasets and annotating the scATAC-seq clusters using scRNA-seq derived labels (**Fig. S2G**). Thus, Tregs, Tconvs, and follicular cells exhibit strong, distinct chromatin and transcriptional identities. In contrast, cell states -such as cycling and interferon response-greatly alter transcriptome profiles without significantly affecting chromatin accessibility profiles.

To identify transcription factors (TFs) that may regulate the gene expression program in each cell cluster, we selected “positive TF-regulators”,^45^ defined as TFs whose gene expression correlates positively with the accessibility of their respective binding motifs across the genome (TF-binding motifs, TFBM). The strongest drivers of cluster-specific molecular programs included *JUN/JUNB, FOS/FOSB, BATF, RORA, RUNX2, EOMES, LEF1, MAF*, and *IRF4* (**Fig. 2D-E** and **S2H**). Of note, the gene expression of certain TFs, such as *FOXP3* and *PRDM1* (encoding BLIMP-1), was negatively correlated with the accessibility of their respective TFBM (i.e., in *FOXP3*-expressing cells, chromatin binding sites for this TF were less accessible). While this may be related to repressor activity, these TFs were excluded from further analysis due to the challenges associated with their interpretation.

Using the identified positive TF-regulators, we generated a heatmap depicting significant TFBM enrichment across all CD4+ T cell subsets, and performed unsupervised clustering (**Fig. 2F**, and **Table 7**) which identified three main TF-driven groups: 1) naïve/memory Tconv and Treg clusters; 2) Tconv effectors; and 3) a group including follicular (Tconv-Tfh, Tconv-CXCL13 and Treg-Tfr) and tissue-resident clusters (Treg-Trm and Tconv-Trm), which exhibited shared follicular and tissue-adaptation features.

The naïve/memory program was regulated by *TCF7* and *LEF1,*^46^ while the Tconv effector phenotype was guided by *KLF2, EOMES, TBX21, RORA*, and *RORC,*^47^alongside *JUN, FOS, REL, HIVEP1-3, NFE2L*, and *MAF* family members^47^, which were also shared by Trm and follicular CD4+ T cells. Meanwhile, follicular and Trm cells showed strong enrichment for *POU2F1-3*, *NFATC2, CIC*, *ARID3A, MAF*, and *BATF.* Interestingly, POU family members^48,48^ and *NFATC2*^49,50^ have been involved in T cell development, activation and differentiation; while *CIC*^51^ plays a role in regulating Tfh cell differentiation and germinal center responses. *ARID3A,*^52^ although linked to autoimmunity, has a unclear role in T cells. Additionally, the TFs MAF and BATF are well-established regulators of follicular CD4+ T cell programs,^53–56^ while also promoting tissue residency in Tregs.^31,55^ BATF has further been associated with Treg stability and function within the TME.^32^ In line with these findings, BATF motif enrichment or TFBM, footprint, and protein expression (mainly in LNs and tumors) were higher in follicular and Treg-Trm than other memory or naïve populations (**Fig. 2G-I** and **S2I**). In addition, the TFBMs of *MAF* and *IRF4*, both previously linked with Tfh phenotypes, were also highly accessible in all follicular and tissue resident subsets (**Fig. S2J**).

Consistent with scRNA-seq and FACS observations, follicular T cell subsets showed strong similarities to tissue-resident cells in both locations (**Fig. 1C-D, 1H** and **2B**), characterized by enrichment of tissue-homing molecules (*CXCR3*, *CXCR6*, *CD69*, *IL1R1*, and *PRDM1*) and reduced expression of circulating markers (*CCR7*, *SELL*, *S1PR1*, and *KLF2*). These findings suggest that all identified follicular subsets (Treg-Tfr, Tconv-Tfh, and Tconv-CXCL13) as well as Treg-Trm and Tconv-Trm share a conserved molecular program integrating TCR activation and tissue homing, closely aligning with the core features of tissue residency.

To further refine this characterization, we constructed the TF:gene regulatory networks^57^ of the follicular and tissue resident clusters (**Fig. 2J**, **Fig. S2K,** and **Table 8**). Beyond shared regulatory mechanisms of T cell activation (*JUN/B/D, FOS/B*) and tissue residency (*BATF*) among Tfh-like and Trm subsets, their TF-target gene architectures were distinctly organized, reinforcing their unique cell identities. Notably, *BATF* was central to the follicular and tissue resident networks and was predicted to regulate distinct gene sets in different T cell types. For example, among Tconvs, *BATF* was predicted to regulate the expression *CD69* and *CD44* in Tconv-Trm, *IFNG, CXCL13* and *TOX2* in Tconv-CXCL13 and/or Tconv-Tfh, the later experimentally validated in the literature.^58^ Conversely, among Tregs, *BATF* was associated with a different set of genes, including *IL21R, SOCS1,* and *NFKB2* in the two Treg populations; *ZNF281* in Treg-Tfr, and *CCR8* and *TNFRSF9* in Treg-Trm.

In summary, combined scATAC-seq and scRNA-seq analyses revealed that follicular CD4+ T cell subsets (Tconv-Tfh, Tconv-CXCL13, and Treg-Tfr) and tissue resident cells (Treg-Trm and Tconv-Trm) share a distinct gene regulatory program that integrates follicular identity, TCR activation, and tissue adaptation. This molecular program not only defines their unique identity but also reinforces their coordinate program in the tumor microenvironment.

### Tumor-Reactive Follicular CD4⁺ T Cells Circulate Between Lymph Nodes and Tumors

Follicular and tissue resident CD4⁺ T cells are frequently observed in tumors and exhibit features consistent with active tumor responses, yet their clonal architecture in invaded LNs and functional contributions to tumor immunity remain poorly understood. To address these gaps, we analyzed their clonal expansion, tumor reactivity, and TCR sharing across LNs and tumors. We integrated transcriptomic data and paired TCRα/β sequences from three patients, obtaining 16,865 cells corresponding to 12,810 different clonotypes from all the clusters and tissues, with no clonotypes shared between patients (**Table 9**).

Expanded clones (defined as two or more cells expressing the same α/β TCR) were localized within distinct UMAP regions, with distribution patterns differing between blood and tissues (**Fig. 3A**). To further investigate the distribution of clonotypes across subsets, we computationally reassigned cells from state-driven clusters to either Treg or Tconv populations based on expression similarities (**Fig. S3A**). We then visualized clonal frequency (pie charts) and absolute numbers (bar plots) by cluster (**Fig. 3B**) and quantified clonal expansion using the Gini index for each patient and tissue (**Fig. 3SB)**. Clonal expansion patterns varied significantly across tissues. In blood, 29% of cells belonged to expanded clones, with the lowest TCR diversity observed in effector Tconvs (mainly Tconv-GZMH) and central memory Tregs. Circulating naïve and quiescent Tregs and Tconv cells displayed the highest diversity. LNs and tumors showed similar expansion patterns, with 31% and 52% of cells deriving from expanded clones, respectively. Expanded Tconv clonotypes in LNs primarily exhibited effector phenotypes (GZMH, GZMK, Trm, and CXCL13), whereas expanded Tregs spanned all non-naïve clusters. In tumors, however, expanded Tregs were more restricted to terminal effectors, effector-memory, Trm, and follicular phenotypes (**Fig. 3C and S3B-C**).

**Fig. 3.**
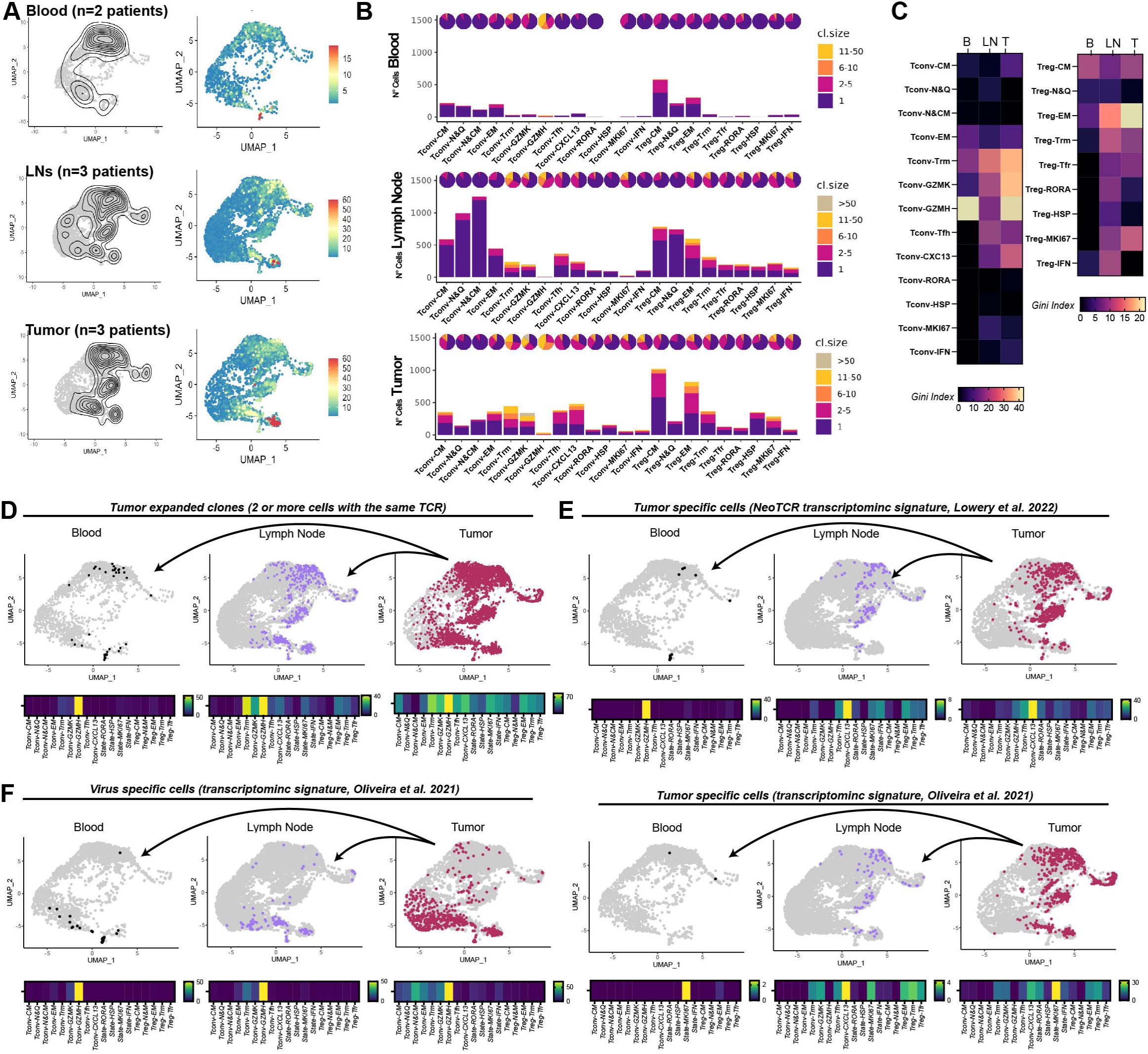
TCR sequencing reveals the phenotypic landscape of clonal expansions across tissues. **(A)** Per-tissue distribution of cells belonging to expanded TCRαβ clones (>1 cell/clone). UMAP projections display individual cells overlaid with a contour plot indicating the density of expanded TCRs (**left panels**) or individual cells colored by clonal size (**right panels**). **(B)** Pie charts (**top**) depict the proportion, and bar plots (**bottom**) show the number of cells per cluster across the three tissues, colored by clonal size. **(C)** Heatmap of the Gini Index for each cluster/tissue. Tumor-expanded clusters are highlighted. **(D-F) Top:** UMAP projections illustrating the distribution of cells belonging to tumor-expanded clones (**D**) or cells with high tumor-specific signature scores (**E-F**) across tissues. **Bottom:** Heatmaps showing the percentage of these cells across clusters by tissue. Applied signatures include Neo-Antigen Specific CD4+ T cells (**E**) from Lowery et al. 2022; Neo-Antigen Specific (**E**) and Virus-Specific CD8+ T cells (**F**) from Oliveira et al. 2022.

We tracked tumor-expanded clones across tissues using TCRs as lineage barcodes. Of the clones seen in more than one tissue, 7% (200 cells across 63 clones) were detected in the blood, primarily within the Tconv-GZMH cluster. Of tumor-expanded clones (2,019 cells across 450 clones) 50% were also identified in lymph nodes, with the highest enrichment observed in the Tconv-GZMH, Tconv-Trm, Tconv-GZMK, and to a lesser extent, Tconv-CXCL13 clusters. Within the Treg compartment, tumor-expanded clones shared with LNs were predominantly localized to the Treg-EM and Treg-Trm clusters (**Fig. 3D**).

To investigate potential tumor reactivity, we analyzed two tumor neoantigen-specific CD4+ T cell transcriptomic signatures.^59,60^ Using the approach developed by Lowery et al.^59^, we identified a total of 735 cells (corresponding to 257 clones) as neoantigen-specific T cells within tumors (**Fig. 3E, upper panels**). This signature was predominantly enriched in Tconv-CXCL13 and, to a lesser extent, in Tconv-Tfh, Tconv-RORA, and Tconv-GZMH, suggesting *in-situ* T cell activation and potential tumor reactivity. In the LNs, these neoantigen-specific clones were distributed across multiple subsets, including the Tconv-CXCL13, Tconv-Tfh, Tconv-Trm, as well as cycling T cells and interferon-response populations. In contrast, in the blood, these clonotypes were primarily confined to the Tconv-GZMH cluster. Notably, a similar distribution pattern was observed using the signature described by Oliveira et al.^60^ (**Fig. 3E, bottom panels**), except for Tconv-GZMH, which showed no significant enrichment. Overall, T cells with neoantigen features were particularly enriched in follicular and proliferative subsets within LNs and tumors. For Tregs, neoantigen signatures were detected in Treg-Trm and Treg-Tfr populations; however, their interpretation remains challenging as these signatures have not been functionally validated for Tregs.^59^

To test these predictions experimentally, we conducted an *in vitro* tumor antigen specificity assay. Total cell suspensions from invaded LNs were cultured to allow tumor-antigen presentation, leading to T cell activation and subsequent 4-1BB upregulation (**Fig. S3D**). This response was blocked by the addition of an anti-human HLA-DR antibody. Among Tconvs, Tconv-Tfh and Tconv-CXCL13 cells exhibited the highest levels of 4-1BB expression, which was reduced by HLA-DR blockade, supporting their tumor antigen reactivity. These assays also suggest tumor reactivity within Treg-Tfr cells. Notably, both Treg-Tfr and Treg-Trm expressed 4-1BB in culture, but only Treg-Tfr showed reduced expression following HLA-DR blockade. However, interpretation for Tregs remains challenging as they display basal 4-1BB and HLA-DR expression.

To contrast tumor reactivity with unrelated antigen specificities, we analyzed a virus-specific signature and cross-referenced detected TCRs with public databases. The virus-specific signature primarily highlighted cells within the Tconv-GZMH cluster, a subset associated with chronic viral infections (**Fig. 3F**), consistent with a previous report ^61^. Among the 3,411 tumor-expanded T cells (spanning 901 clones), only 84 cells (23 clones) expressed autoimmunity-associated TCRs, and 74 cells (24 clones) expressed known pathogen-associated TCRs, with no distinct patterns observed (**Fig. S3E**).

Overall, expanded CD4⁺ T cell subsets, including both Tregs and Tconvs, traffic between invaded LNs and tumors, highlighting the trafficking of potential tumor-reactive clones through the LN–tumor axis. Expanded Tregs were enriched in Trm and effector-memory clusters and, together with Treg-Tfr, exhibited neoantigen-associated signatures, although their antigen specificity remains difficult to interpret. Notably, Tconv subsets (including GZMK, Trm, and follicular-like cells) were shared across LNs and tumors, with Tconv-CXCL13 consistently showing neoantigen signatures and in vitro tumor reactivity.

### Follicular and Treg-Trm CD4⁺ T Cells Share Overlapping TCR Repertoires across Lymph Nodes and Tumors

Given their close similarity, we sought to analyze the relationships between follicular and Treg-Trm CD4⁺ T cells by examining TCR sharing, which reflects not only clonal expansion but also transitions in cell state.^62^ To this end, we analyzed the TCR overlap within our scRNA-seq dataset, using naïve and quiescent CD4⁺ T cells as reference populations.

We first visualized repertoire intersections using an Upset plot (**Fig. 4A, left**), which depicts the predominant shared clones across subsets and the total number of clones involved in each sharing (shown at the top of the graphic). Our analysis revealed that Tconv-Tfh and Tconv-CXCL13 subsets shared 94 clones, whereas Treg-Tfr shared 16 clones with Treg-Trm. Notably, in the context of potential Treg–Tconv transitions (illustrated in violet), despite having fewer overall clonotypes, Treg-Tfr exhibited TCR sharing with Tconv-CXCL13 (7 clones) and with Tconv-Tfh (5 clones). As expected, the high TCR diversity in naïve and quiescent (N&Q) Tconv and Treg cells led to some clonotype sharing with follicular populations and with each other; however, significant overlap beyond random expectation was observed specifically between Tconv-Tfh and Tconv-CXCL13, as well as between Tconv-Tfh and Treg-Tfr, as determined by Morisita-Horn index analysis (statistics in **Fig. 4A**, **Table 9** and **Methods**). Circos plots (**Fig. 4A, right**) further illustrated substantial TCR repertoire overlap between Tfh and CXCL13 subsets, Treg-Tfr and Treg-Trm clusters, and between Treg-Tfr and both CXCL13 and Tfh populations.

**Fig. 4.**
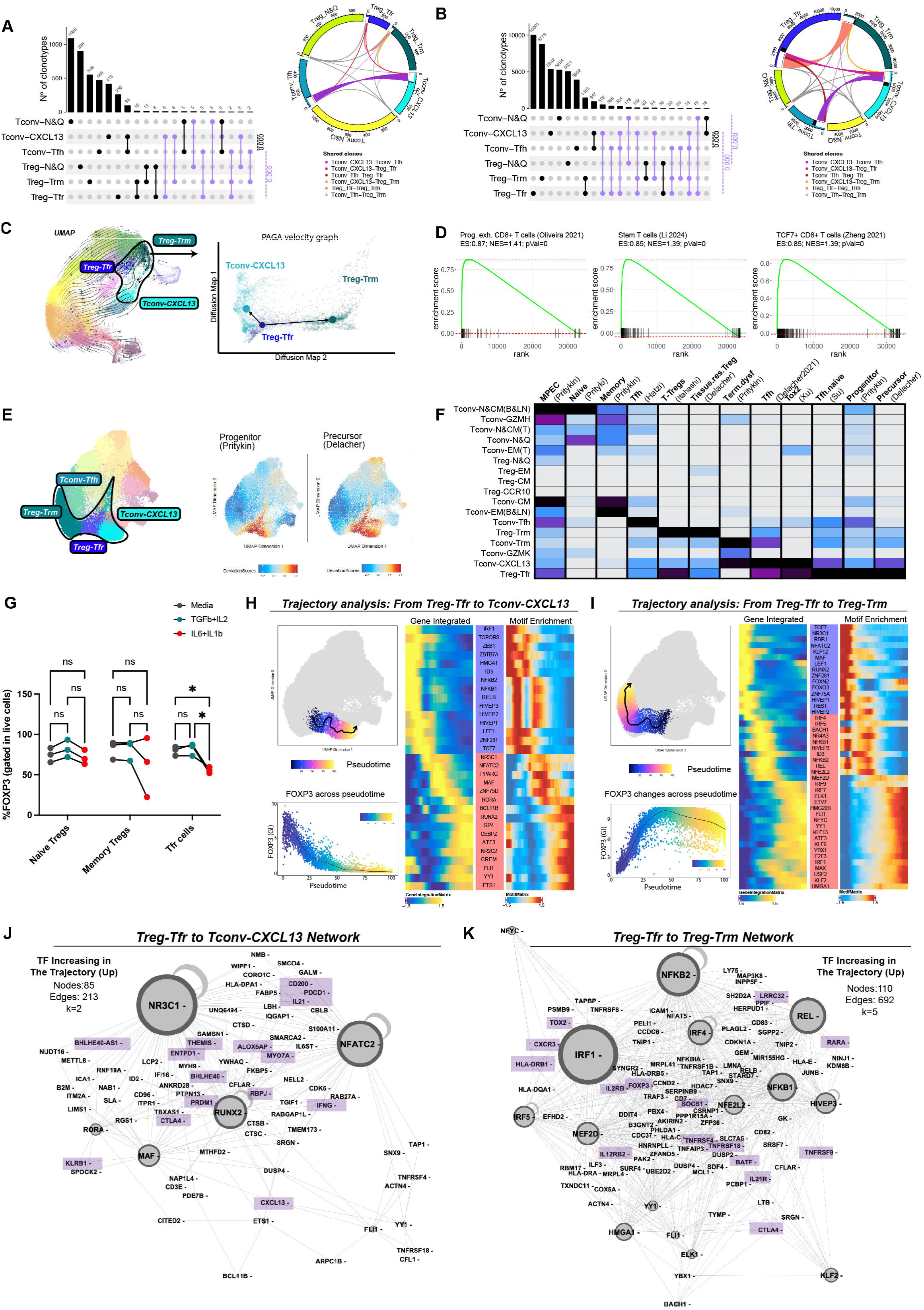
Integrated analyses support Treg-Tfr dual plasticity to Tconv-CXCL13 and Treg-Trm. **(A-B)** TCR repertoire overlap across follicular and naïve Tregs and Tconv subpopulations. (**A)** Single-cell TCR-seq analysis; **(B)** Bulk RNA-seq analysis. **(A-B, left)** Upset plots depicting the number of clones/cells at each intersection. Lines and circles indicate the populations involved in each comparison. Violet lines denote the Treg-Tconv sharing patterns. Statistical significance was assessed relative to Morisita-Horn Indexes for each comparison and random expectation within each comparison (p-value calculated via permutation; see Methods). **(A-B, right)** Circos plots visualizing clonotype distribution across populations. Clonotypes shared between two subsets are highlighted in **white**, while those shared across multiple subsets appear in **black**. Key sharing patterns are color-coded. **(C)** scRNA-seq Velocity analysis: UMAP plot (left) and Diffusion map plot (right) showing RNA velocity vectors and PAGA projection of cells from lymph node and tumor cells (scRNA-seq dataset). Streamlines depict directional flow for all clusters (stochastic model, **right**) and follicular subsets (stochastic model, **left**), with line weight indicating velocity. **D)** Gene set enrichment analysis (GSEA) plots showing the Normalized Enrichment Scores (NES) and p-palues of Treg-Tfr cluster for publicly available signatures of progenitor exhausted CD8+ T cells and Stem CD4+ T cells. **(E-F)** scATAC-seq analysis: Peak-signature enrichment public signatures for follicular cells, progenitor/precursor cells, tissue-Treg cells, dysfunctional cells, and exhausted cells **(E)** UMAP projection of per-cell peak signature enrichment. **(F)** Heatmap displaying adjusted p-values of normalized pseudo-bulk enrichment. (**G)** In vitro polarization assay: Treg-Tfr, Treg-Naive and Treg-Trm were FACS-sorted from healthy donor PBMCs and stimulated 3 days with anti-CD3/CD28 under different cytokine conditions as indicated. Scatter plots show the percentage of FOXP3+ cells within live cells. Statistical test: Two-way ANOVA (paired) with Tukey’s multiple comparisons post hoc test (* = p ≤ 0.0332). **(H-I)** ATAC-seq trajectory analysis. **(H)** Transition from Treg-Tfr to Tconv-CXCL13. **(I)** Transition from Treg-Tfr to Treg-Trm. **Upper left**: UMAP visualization of pseudo-time ordering of nuclei along differentiation trajectories**. Botton left** each UMAP, integrated gene expression of *FOXP3* across pseudo-time is shown. **Right panels** display heatmaps of correlated gene integration (GI) and transcription factor (TF) motif enrichment along the trajectory. **(J-K)** TF-target gene networks: Regulatory networks driving Treg-Tfr differentiation to Tconv-CXCL13 (J) and Treg-Trm. Node size represents out-degree, with selected target genes highlighted in squares.

To increase statistical power in evaluating TCR repertoire overlaps and to validate in additional patients, we performed bulk TCR-seq on the same samples sorted for the bulk-RNA dataset (**Fig. 4B** and **Table 10**). This strategy increased the number of detected clones in each group by an order of magnitude, particularly improving sensitivity for the Treg-Tfr subset. Both UpSet and Circos plots confirmed substantial TCR sharing between Tconv-Tfh and Tconv-CXCL13 and highlighted notable overlap of Treg-Tfr with Treg-Trm and Tconv-CXCL13. Statistically significant overlap was observed specifically between Tconv-Tfh and Tconv-CXCL13, as well as between Treg-Tfr and Tconv-CXCL13 and Treg-Tfr and Tconv-Tfh.

Overall, our findings reveal substantial TCR repertoire sharing among follicular-like CD4⁺ T cell subsets, including Treg-Trm cells. This overlap underscores the plasticity of these populations, particularly Treg-Tfr, demonstrating their capacity to transition between functional states in response to tumor-associated cues. Such adaptability highlights the role of Treg-Tfr and other follicular-like cells in shaping anti-tumor immune responses and their potential as immunotherapeutic targets.

### Treg-Tfr can differentiate to both Treg-Trm and ex-Tregs adopting features of Tconv-CXCL13 cells

To elucidate the developmental relationships of Treg-Tfr cells with Treg-Trm and Tconv-CXCL13 populations, we performed RNA velocity analysis to infer differentiation trajectories. Velocity vectors across the full UMAP confirmed expected differentiation pathways from naïve and quiescent cells toward effector states in both conventional and regulatory populations, with prominent transitions involving follicular-like subsets (**Fig. 4C, left**). Interestingly, when focusing specifically on Treg-Tfr relationships, predicted velocity vectors suggested bidirectional differentiation potential from Treg-Tfr toward both Tconv-CXCL13 and Treg-Trm populations (**Fig. 4C, right**). Additionally, analysis of conventional follicular transitions revealed that the predominant pathway proceeds from Tconv-Tfh to Tconv-CXCL13 (**Fig. S4A**), supporting distinct developmental routes for follicular cell generation.

To further investigate the progenitor potential of Treg-Tfr cells suggested by velocity analysis, we performed comprehensive molecular and epigenetic characterization. First, we confirmed the purity and identity of Treg-Tfr populations through multiple validation approaches (**Fig. S4B-D**). These cells exhibited canonical Tfr signatures (**Fig. S4B**), expressed FOXP3 protein by flow cytometry (**Fig. S4C**), and showed high prediction scores for regulatory identity when analyzed using transfer learning algorithms (**Fig. S4D**). We then specifically assessed their progenitor status by comparing Treg-Tfr cells with established progenitor populations and contrasting them with other follicular-like subsets exhibiting more differentiated characteristics, thereby validating our inferred trajectories. Chromatin accessibility profiling revealed that Treg-Tfr cells shared significant ATAC-seq peak profiles with multiple stem-like signatures, including progenitor exhausted T cells^63,64^ and CD4+ T stem cell populations^55^ (**Fig. 4E-F** and **Table 11**). Notably, Treg-Tfr cells exhibited accessibility patterns consistent with tissue-Treg precursors previously identified in LNs, also characterized by BATF transcriptional programming.^55^ Uniquely, Treg-Tfr cells within the tumor microenvironment (invaded LNs and tumors) expressed distinct molecular markers including COCH, ICA1, PMC, and ZNF281 (**Fig. S4F-H**), suggesting tissue-specific adaptations while maintaining progenitor characteristics. In contrast, Tconv-CXCL13 clusters displayed epigenetic signatures consistent with tumor-infiltrating Tfh and exhausted/dysfunctional states^64,65^, while Treg-Trm showed canonical tumor-Treg features^31,66^ (**Fig. 4E** and **S4E**).

Supporting the hypothesis of ex-Treg development, lineage-tracing studies in mice have demonstrated that 10-20% of Treg-Tfr cells lose FOXP3 under inflammatory conditions, differentiating into follicular Tconv cells.^67,68^ Furthermore, Tconv-CXCL13 cells chromatin accessibility at the *FOXP3, RORC*, and *IL17A/F* loci resembled the epigenetic signature of murine ex-Tregs, characterized by the acquisition of an Th17-like inflammatory program (**Fig. S4I**), as reported in autoimmunity.^69^ Consistently, the transcriptome (**Fig. S4J)** and epigenome (**Fig. S4K**) of Tconv-CXCL13 was enriched in Th17 related genes. Functionally, flow cytometric analysis across all three patients revealed that Tconv-CXCL13 cells produced high levels of IL-21 and IFN-γ, either individually or in combination, consistent with their follicular helper and inflammatory characteristics (**Fig. S4L)**.

To directly test the plasticity of Treg-Tfr cells, we performed *in vitro* differentiation assays using cells isolated from healthy donor PBMCs, from which Treg-Tfr cells can be obtained in sufficient quantities for functional analysis. Under conventional T cell-polarizing conditions including IL-6 and IL-1ß,^70,71^ Treg-Tfr cells exhibited significantly greater propensity to lose FOXP3 expression compared to memory, effector, and naïve Treg counterparts (**Fig. 4G** and **Fig. S4M**). These findings support a model wherein Treg-to-conventional cell transitions occur preferentially through Treg-Tfr intermediates, potentially generating Tconv-CXCL13 cells with combined follicular helper and inflammatory Th17-like characteristics.

Overall, our data demonstrate that Treg-Tfr cells undergo plastic conversion within tumors. These cells exhibit multipotent progenitor signatures and retain bidirectional differentiation capacity toward regulatory (Treg-Trm) lineages, but, notably can also transdifferentiate into ex-Tregs with Tconv-CXCL13 features. Collectively, these findings identify Treg-Tfr cells as key regulators of the effector-regulatory balance in tumor follicular responses, providing insights for their therapeutic targeting to enhance anti-tumor immunity.

### Progenitor-like Treg-Tfr Bifurcation Through Alternative Gene Regulatory Programs

Given the observed plasticity of Treg-Tfr cells, we next sought to uncover the transcriptional regulators governing their alternative differentiation pathways. We investigated the TF regulatory programs underlying Treg-Tfr developmental trajectories: maintenance of regulatory identity (Treg-Tfr to Treg-Trm) versus conversion to effector cells (Treg-Tfr to Tconv-CXCL13). As a comparative control, we also analyzed the Tconv-Tfh to Tconv-CXCL13 transition to distinguish Treg-to-Tconv conversion mechanisms from conventional Tconv developmental programs. Using scATAC-seq data, we performed trajectory analysis through pseudotime ordering of TFs whose motif enrichment in differential peaks positively correlated with TF expression (gene score and integrated gene expression). The resulting gene expression and motif enrichment pseudotime heatmaps (**Fig. 4H-I**) revealed distinct TF dynamics driving these transitions toward terminally differentiated regulatory and effector cells.

Analysis of Treg-Tfr trajectories revealed opposing TF activity patterns between identity maintenance and conversion to effector cells. During the Treg-Tfr to Treg-Trm transition, we observed upregulation of ID3, NFKB1-2, HIVEP3, HMGA1, and IRF1 (a transcriptional regulator of IFNs and IFN-inducible genes involved in Tr1 differentiation^72^), alongside unique increases in NFE2L2, IRF4, REL, ETV7, and MEF2D—transcriptional regulators controlling suppressive function and expression of IL-10, CTLA-4, and ICOS in Treg cells.^73^ Conversely, NFATC2, MAF, RUNX2, and NR3C1 (repressor of effector T cells exhaustion)^74^ decreased along this trajectory.

In contrast, the conversion pathway from Treg-Tfr to Tconv-CXCL13 displayed a complete reversal of this regulatory program. ID3, NFKB1-2, HIVEP3, HMGA1, and IRF1 were progressively lost, while NFATC2, MAF, RUNX2, and NR3C1 increased. This Treg-to-Tconv conversion uniquely upregulated PPARG, CREM, and RORA (regulators of Th17 gene signatures),^75^ as well as ATF3, YY1, NR2C2, BCL11B, and FLI1 (inflammatory mediators).^76–82^

To distinguish Treg-specific conversion from general Tconv maturation, we compared both trajectories converging on Tconv-CXCL13 (**Fig. S4N**). Both transitions shared downregulation of stemness-associated factors LEF1 and TCF7^83^, alongside upregulation of RORA, MAF, RUNX2, PPARG, and NFATC2, which are linked to Th17 differentiation, tissue residency, and T cell activation.^84–88^ However, key differences distinguished Treg conversion from Tconv maturation: ETS1, which mediates T cell activation, Th17 responses, and autoimmunity,^89^ was upregulated during Treg-Tfr to Tconv-CXCL13 conversion but downregulated during Tconv-Tfh maturation. Conversely, RELB, HIVEP3, and NFKB2 (key regulators of T cell and iNKT activation),^90–92^ showed the opposite pattern. The control Tconv-Tfh to Tconv-CXCL13 transition specifically upregulated BATF, RUNX3, TBX21, JUN, NR4A1, IRF5, and IRF9 (regulators of tissue residency, TCR activation, and IFN responses), ^93–98^ features absent in Treg conversion.

To define mechanistic regulatory relationships, we constructed TF-target networks by integrating trajectory-enriched TFs with differentially expressed genes (DEGs) containing differentially accessible cognate TF binding motifs in terminal cell populations (**Fig. 4J-K and S4N**). These networks revealed distinct regulatory patterns for Treg-Tfr fate decisions. During identity maintenance (Treg-Tfr to Treg-Trm), IRF1 emerged as the primary regulator, maintaining expression of key Treg genes including FOXP3, CTLA4, LRRC32 (GARP), and RARA. During conversion to effector cells (Treg-Tfr to Tconv-CXCL13), NR3C1 emerged as the dominant regulator, linking for the largest proportion of Tconv-CXCL13 genes. NR3C1 cooperated with NFATC2, RUNX2, RORA, and MAF to drive genes associated with Tconv identity (BHLHE40, THEMIS) and effector function (IL21, IFNG, CXCL13). Notably, these master regulators of Treg-Tfr plasticity, IRF1 for identity maintenance and NR3C1 for conversion, respond to distinct microenvironmental cues (IFN cytokines and glucocorticoids, respectively), suggesting that Treg-Tfr fate is environmentally regulated.^65,99,100^

In contrast, the control Tconv-Tfh to Tconv-CXCL13 transition revealed BATF as the key driver, collaborating with JUN, NR4A1, NFKB2, REL, and RELB. This pattern suggests enhancement of Tconv follicular and activation programs rather than dramatic reprogramming, highlighting the unique nature of Treg-to-Tconv conversion.

Collectively, our analysis revealed that the two alternative fates for Treg-Tfr cells, maintenance of regulatory identity or conversion to ex-Tregs, are controlled by distinct transcriptional programs: among other TFs, IRF1 likely drives regulatory while NR3C1 orchestrates effector conversion. These findings identify Treg-Tfr cells as potential sensors of microenvironmental cues and provide specific transcriptional targets for therapeutic manipulation of CD4+ T cell fates.

### Follicular T Cells Maintain Identity While Adapting to Tissue Microenvironments During Migration Between Invaded Lymph Nodes and Tumors

TCR sharing between cells informs not only on clonal expansion and state transitions, but also on cell migration patterns when identical TCRs are found across different tissues. To investigate T cell trafficking in our dataset, we analyzed migration of Tregs and Tconvs between tissues at the population level, pooling all pure and mixed clusters. Using the normalized Morisita-Horn index (MHI: 0 = no sharing, 1 = complete sharing), we found that both Tregs and Tconvs migrated primarily between LNs and tumors (Tregs: MHI = 0.19, 299 shared clones; Tconvs: MHI = 0.18, 329 shared clones), with minimal trafficking to blood (**Fig. 5A**).

**Fig. 5.**
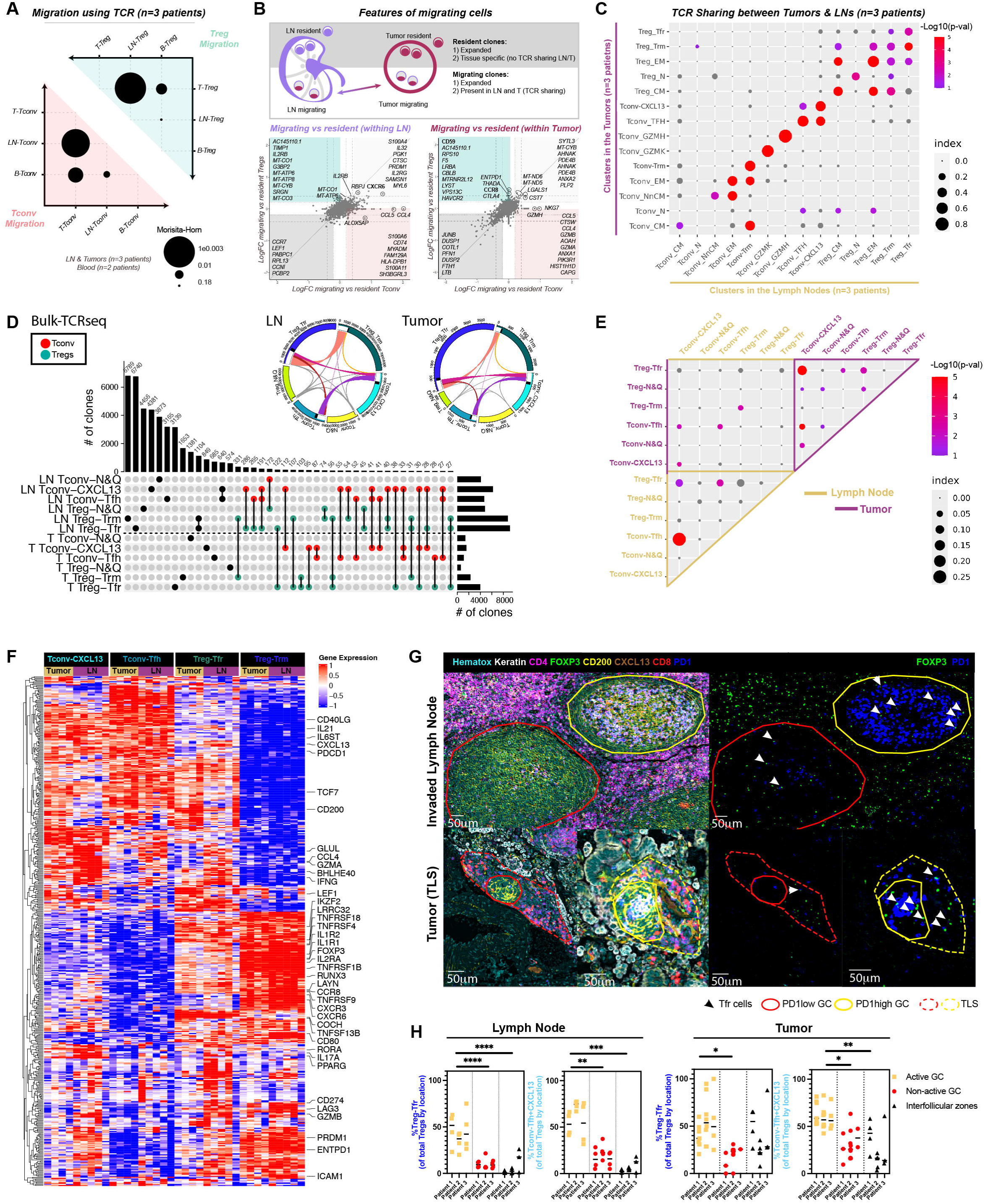
TCR sharing analysis reveal patterns of migration (LN-tumor network) and transcriptomic and spatial analysis reveal LN-Tumor similar profile and location of Tfh-like subsets. (**A)** Scatter plots visualizing Treg/Tconv shared TCRaβ clones. Dot size is proportional to the Normalized Morisita-Horn Index (MHI) of intersected groups, calculated either among total Tregs (green triangle), or Tconvs (red triangle) among blood, lymph node or tumor. (**B) Upper panels**: Scheme illustrating the definition of resident and migrating clones. **Bottom panels**: Log2 FC/FC plots comparing migrating and resident cells per tissue (Log2FC>0.2 and adj-pVal<0.05). Selected genes from each comparison are displayed in the corresponding squares. (**C)** General LN-tumor intersection: Dot plot visualizing the Morisita-Horn Indexes (size of the circle) measuring pairwise TCR repertoire overlap between LN and tumor T cell populations or tissues. The color of circles indicates significance compared to random expectation (p-value calculated by permutation). **(D)** Bulk TCR-seq analysis. TCR repertoire overlap across Treg-Trm, follicular and naïve Tregs and Tconv subpopulations split by tissue. Upset plot display the number of clones/cells at each intersection. Lines (black) and circles (red for Tconvs and green for Tregs) indicate the populations involved in each comparison. Dot-line help to visualize the separation between LNs and tumors. Circos plots visualize clonotype distribution across populations in each tissue. Clonotypes shared between two subsets are highlighted in white, while those shared across multiple subsets appear in black. **(E)** LN-Tumor intersection with focus in follicular and Treg-Trem cells. Dot plot visualizing the Morisita-Horn Indexes (size of the circle) measuring pairwise TCR repertoire overlap between LN and tumor T cell populations or tissues. The color of circles indicates significance compared to random expectation (p-value calculated by permutation). **(F)** Heatmap highlighting the top 500 most deferentially expressed genes (MVGs) characterizing follicular populations in LNs and tumors. **(G-H)** Multiplex immunohistochemistry images of LNs and tumors. **(G)** Representative Images of LNs (upper panels) and tumors (bottom panels) showing antibody combinations as indicated. Red and yellow zones indicate non-active (PD1⁺low) and active (PD1⁺high) germinal centers (GCs), respectively. **(H)** Bar plots quantifying the per-tissue percentage of follicular Treg (CD4⁺ FOXP3⁺CD200⁺PD1⁺) and follicular Tconv (CD4⁺ FOXP3⁻CD200⁺PD1⁺) cells among total Tregs or Tconvs in distinct regions: active GCs (PD1⁺high GC), non-active GCs (PD1⁺low GC), and interfollicular zones. Statistical analysis: paired nested one-way ANOVA with Tukey’s multiple comparisons post hoc test (* = p ≤ 0.0332; ** = p ≤ 0.0021; *** = p ≤ 0.0002; **** = p ≤ 0.0001). Immunohistochemistry analysis: n = 3 LNs and Tumors from NSCLC patients.

To characterize how migrating cells adapt while maintaining identity, we performed differential gene expression analysis comparing migrating and resident populations within each lineage, followed by Ingenuity Pathway Analysis (IPA, **Fig. 5B, S5A**, and **Table S11**). Migrating tumor Tregs uniquely upregulated CCR8^29^, CD59^101^ and ENTPD1 (encoding CD39), which represent potential therapeutic targets for selectively modulating Treg migration and suppressive function.

Cluster-specific analysis of migrating cells, excluding state-driven clusters to avoid in silico re-assignment artifacts, revealed that most clones maintained their transcriptional identity while trafficking between LNs and tumors (diagonal circles in **Fig. 5C**). State transitions occurred primarily among Treg subsets, suggesting high environmental adaptability while preserving core regulatory programs.

To specifically examine follicular and Treg-Trm subsets dynamics across tissues, we generated UpSet and Morisita-Horn index plots comparing LN and tumor populations. Follicular subsets showed substantial TCR sharing within each tissue, particularly Tconv-Tfh and Treg-Tfr with Tconv-CXCL13 (**Fig. 5D-E** and **S5B-C**). Importantly, cells largely preserved their follicular identity during LN-tumor migration.

To examine tissue-specific adaptation, we leveraged our bulk RNA-seq data, which provides deeper sequencing coverage than single-cell. Analysis of the top 500 variable genes revealed that cell population identity remained the primary driver of gene expression, though distinct tissue-specific changes emerged that varied by cell population (**Fig. 5F** and **S5D**). Treg-Tfr showed minimal tissue-dependent variation (10 LN-upregulated, 7 tumor-upregulated genes), maintaining remarkably stable identity across environments. In contrast, Tconv-Tfh exhibited the greatest tissue-specific adaptation (177 LN-upregulated, 51 tumor-upregulated genes) while preserving core follicular characteristics. Treg-Trm and Tconv-CXCL13 displayed intermediate adaptation patterns (Treg-Trm: 39 LN-upregulated, 33 tumor-upregulated; Tconv-CXCL13: 60 LN-upregulated, 29 tumor-upregulated).

Across T cell subsets, transcriptional differences between lymph node and tumor compartments were moderate but consistent. LN populations showed higher expression of genes linked to migration, metabolic flexibility, and immune regulation, such as S1PR1, PDGFD, MAOB, LAMB2, and SLCO2B1, consistent with trafficking, stromal responsiveness, and maintenance of regulatory programs. In contrast, Tumor-resident cells upregulated genes associated with tissue retention, adhesion, and effector function, including EPCAM, MSLN, CEACAM6, CD24, MAL2, CXCR2, CD69, TNFRSF4, and XIRP1, reflecting enhanced stromal anchoring, chemotaxis, and activation. Subset-specific patterns emerged: Treg-Trm and Treg-Tfr retained stronger regulatory/metabolic signatures in LNs, whereas Tconv-Tfh and Tconv-CXCL13 exhibited moderate expression of effector and adhesion modules in tumors.

To determine whether spatial organization reflects this balance of identity and adaptation, we performed multiplex immunohistochemistry on NSCLC patient samples. Follicular populations maintained their germinal center tropism across tissues: in LNs, Tconv-Tfh and Tconv-CXCL13 (PD1+CD200+ Tconvs) and Treg-Tfr (PD1+CD200+ Tregs) concentrated in mature germinal centers (PD1high) versus immature GCs or interfollicular zones (**Fig. 5G-H**). This spatial preference persisted in tumors, where these populations localized to mature tertiary lymphoid structures (TLS), specifically within germinal centers (**Fig. 5G-H**). Spatial transcriptomic analysis confirmed that all follicular clusters maintained their co-localization patterns within tumor TLS, alongside B cells, preserving lymphoid organization while adapting to the tumor (**Fig. S5E**).

Collectively, our integrated analysis reveals that follicular T cells successfully balance identity maintenance with tissue-specific adaptation during migration between invaded lymph nodes and tumors. Despite both tissues containing cancer cells and sharing TCR repertoires and spatial organization, each tissue imprints distinct transcriptional adaptations on resident populations. The preferential localization of follicular subsets within mature germinal centers and TLS, combined with their tissue-specific transcriptional programs, positions these populations as key orchestrators of local adaptive immune responses. This balance between maintaining core identity while adapting to microenvironmental cues offers multiple therapeutic intervention points for modulating anti-tumor immunity.

## DISCUSSION

Our integrated multi-omic approach reveals the diversity of CD4+ T cell populations within tumor-invaded lymph nodes, demonstrating that these sites function as both sources and maintenance hubs for progenitor, naïve, and specialized populations with tumor-specific characteristics. By combining scRNA-seq, scTCR-seq, and scATAC-seq, we achieved high-resolution characterization through several methodological innovations: equal admixture of Tregs and Tconvs to enhance rare population detection, patient-matched specimens across tissues, and a novel surface marker strategy for FACS-sorting (avoiding reliance on CXCR5 for Tfr cells which is unstable after enzymatic digestion).^102^

Rather than merely facilitating metastatic spread, invaded lymph nodes function as active sites for immune activation and differentiation of tumor-reactive CD4+ T cells. Consistent with CD8+ T cell studies and murine models,^7,8,31,103^ we demonstrate enrichment for CD4+ T cells with anti-tumor potential, including progenitor populations and functionally competent cells bearing both follicular and tissue-resident characteristics. The selective presence of Treg-Tfr, Tconv-Tfh, Tconv-CXCL13, and Treg-Trm within both invaded lymph nodes and tumors, while virtually absent from peripheral blood, suggests specialized niches harboring tumor-specific cells. These populations exhibit tissue residency features, expressing retention markers (CXCR3, CXCR6, CD69, IL1R1, PRDM1) while downregulating circulation determinants (CCR7, SELL, S1PR1, KLF2)^26,104–108^, and show enrichment for tumor antigen-specific signatures, aligning with murine models^109^ and human studies.^110,111^ Moreover, these follicular and tissue-resident subsets are enriched in cells bearing tumor-reactive signatures and upregulate 4-1BB in an HLA-DR dependent manner when total lymph node cell suspensions are cultured. These findings indicate that invaded lymph nodes act as active reservoirs and differentiation hubs for tumor-specific immunity.

Migration analysis using the TCR as a molecular tag identified substantial clonotype sharing between lymph node and tumor populations, providing compelling evidence for trafficking of follicular T cells, complementing previous studies.^6–10^ Notably, migrating cells largely retain their lineage-defining transcriptional programs and germinal center– homing capacities. At the same time, they adapt to tissue-specific microenvironmental contexts through modulation of genes associated with migration, metabolism, and immune regulation, consistent with observations reported in lower-resolution studies.^9,112^ This creates an immunological circuit wherein lymph nodes continuously supply differentiated effector cells while receiving feedback from the tumor microenvironment.Our chromatin accessibility analysis identifies BATF as a central transcriptional regulator coordinating both follicular and tissue-residency CD4+ T cell programs, extending previous observations of its role in tissue residency and tumor Treg biology^66,95,113,114^ We describe that BATF is enriched not only in tumor-associated Treg subsets (both Treg-Tfr and Treg-Trm) but also in follicular Tconv populations, revealing its broad involvement in shaping the follicular CD4+ landscape. The convergence of these distinct programs under BATF regulation suggests a unifying transcriptional mechanism that enables CD4+ populations to maintain their core identity while retaining the capacity for subset interconversion in response to microenvironmental cues. This BATF-centered regulatory architecture appears to govern the activation states, differentiation trajectories, and dynamic plasticity observed among follicular-like subsets within the tumor microenvironment.

Most significantly, integrated lineage analysis identifies Treg-Tfr cells as multipotent progenitors functioning as a critical developmental checkpoint in CD4+ T cell fate determination. RNA velocity, chromatin accessibility, and TCR sharing analyses converge on a model where Treg-Tfr cells possess bidirectional differentiation capacity: either maintaining regulatory identity through differentiation into Treg-Trm cells or undergoing conversion into inflammatory Tconv-CXCL13 effector cells. This discovery aligns with recent findings on progenitor properties in CD8+ and CD4+ Tfh cells,^115–118^ and extends murine observations of CD4+ T cells with stem-cell properties differentiating toward either anti-tumor effector or suppressive phenotypes.^103,109,114,119^ Our results provide the first identification in humans that Treg-Tfr cells exhibit such precursor properties.

The clinical relevance of this bifurcation is underscored by contrasting functional outcomes: Treg-Tfr cells and Treg-Trm inhibit anti-tumor immunity,^120,121^ while Tconv-CXCL13 associate with tertiary lymphoid structure formation and positive anti-PD-1 responses.^22,23^ All three populations express inhibitory checkpoints (PD-1, CTLA-4) but lack exhaustion markers (EOMES, LAG3),^122–124^ which may explain their association with immunotherapy outcomes: While Treg-Tfr have been shown to inhibit anti-tumor immunity during PD1 blockade;^120^ CD8+ CXCR5+ and CD4+ CXCL13+ T cells (resembling Tconv-CXCL13), have been associated with positive responses to anti-PD1.^22,23^ Moreover, TLS formation and CXCL13 expression are now recognized as key biomarkers of response to a-PD1 therapy.^21,125–127^ Supporting this, Tconv-CXCL13 cells in tumors preserve the production of cytokines (IL-21 and IFN-γ) that sustain CD8+ T cell responses.^128,129^

We provide the first evidence of FOXP3 instability in human cancer Tfr cells, extending murine observations of Treg fragility.^67,68^ While typically detrimental in autoimmunity, this instability offers therapeutic opportunities in cancer.^130^ In these lines, we have recently demonstrated that depleting CD74, a molecule associated with tumor-enriched Treg cells, leads to FoxP3 loss and acquisition of effector functions, enhancing tumor rejection in humanized models^131^. Furthermore, our trajectory analysis reveals distinct transcriptional programs governing fate decisions: IRF1-driven pathways appear to maintain regulatory identity, while NR3C1-orchestrated networks drive effector conversion, providing preliminary cues for therapeutic manipulation.^65,99,100^ Whether Tconv-CXCL13 cells derived from different precursors (Tfr and Tfh) exhibit distinct behaviors may explain disparities in their roles across cancer types^20,132^ and warrants further investigation.

CD4+ T cells in invaded lymph nodes share expanded clonotypes with tumor-infiltrating populations while exhibiting fewer exhaustion markers than their tumor counterparts.^103,109,133,134^ The Treg-Tfr bifurcating trajectory opens potential intervention avenues: direct targeting could reduce regulatory cell generation while promoting anti-tumor effector responses. Preserving invaded lymph nodes during immunotherapy, radiotherapy, or chemotherapy treatment before tumor rejection could maintain reservoirs of tumor-reactive CD4+ T cells with stem-like properties, as these tissues function as integrated immunological units where interventions in one compartment influence the other. The tissue-adapted characteristics we identify could also help improving adoptive therapies by engineering CAR-T cells or tumor-infiltrating lymphocytes with follicular and tissue residency programs,^135,136 131 137^

Several limitations warrant consideration. The generalizability across cancer types requires validation,, as does the functional significance of patient heterogeneity in Treg-Tfr plasticity. Future studies should prioritize direct TCR-antigen interaction demonstration, functional validation of conversion impact on clinical outcomes, TFs and network validation, and identification of environmental factors determining Treg-Tfr fate decisions.

Our analysis redefines invaded lymph nodes as active immunological partners with tumors, maintaining specialized CD4+ T cell populations through BATF-coordinated tissue-adapted programs. The discovery of Treg-Tfr bifurcation reveals a critical developmental checkpoint with therapeutic potential for enhancing anti-tumor immunity. By demonstrating regulatory T cell plasticity and its integration with tissue-adapted immunity, our work provides a new framework for understanding CD4+ T cell biology in cancer and identifies intervention points for manipulating the effector-regulatory balance toward improved immunotherapy outcomes.

## MATERIAL AND METHODS

### Clinical sample collection

Matched samples of blood, tumor-proximal LNs and tumors were collected from patients with NSCLC having undergone standard-of-care surgical resection at the Institut Mutualiste Mountsouris (Paris, France). Tissue samples were taking from surgical residues available after histopathological analysis and not required for diagnosis. The human experimental procedures follow the Declaration of Helsinki guidelines and were approved by the Institutional Review Board and Ethics Committee of Institut Curie Hospital group (CRI-DATA190154). All patients signed a written consent. Samples were characterized by IHC, NGS and detection of genomic abnormalities by Cytoscan. The clinical and pathological characteristics of the patients are summarized in **Table 1**.

### Tissue dissociation and cell isolation

Samples were processed within 4 hours after the primary surgery. Tissues were cut into small fragments and digested with 0.1 mg/ml Liberase TL (Roche) in the presence of 0.1 mg/ml DNase (Roche) for 30 min before the addition of CO2 independent medium (GIBCO). Cells were then filtered and mechanically dissociated with a 2,5mL syringe’s plunger on a 40-μm cell strainer (BD) and washed with CO2 independent medium (GIBCO) 0,4% human BSA. Peripheral blood mononuclear cells (PBMCs) were obtained by gradient centrifugation lymphoprep (STEMCELL - Cat#07851), following manufacture instruction.

### FACS-sorting for scRNA-seq and scATAC-seq

For each tissue, Tregs (DAPI-CD45+ CD4+ CD25hi CD127lo) and Tconvs (DAPI-CD45+ CD4+ CD25lo CD127lo/hi) were FACS-sorted and admixed at a fifty/fifty ratio before loading on a 10X Chromium (10X Genomics). Treg and Tconv populations were sorted from fresh samples collected from three different tissues: Blood (B), lymph nodes (LNs) and tumors (T). Tissue dissociation and PBMCs isolation was performed as specified before (Tissue dissociation and cell isolation). Cell suspensions were then washed in PBS and incubated with LIVE/DEAD Fixable Cell Dead Stain (eBioscience) during 15 min at RT. Next, cells were washed, resuspended to a concentration of 30.10^6^ Cells/mL, and incubated with fluorochrome labeled Abs for 30 min at 4°C. Finally, the cells were washed and resuspend in cold MACS buffer to a concentration of 30.10^6^ Cells/mL. Cell-sorting was performed using ARIA Fusion cell sorter (BD). Cells were received in coated Eppendorf (with pure bovine serum for at least 30 minutes at 37°C) containing 350uL of single cell 10x genomics recommended buffer (PBS 0,04% human serum albumin). During the experiment, cells were kept in MACS buffer at 4°C and directly collected into 1,5mL Eppendorf containing PBS 0,04% human serum albumin (YDRALBUM).

### scRNA-seq and scTCR-seq experiment

#### scRNA-seq and scTCR-seq Experiment

10X Genomics Protocols: For the first two patients and the blood sample from the third patient, libraries were prepared using the Single Cell 3′ Reagent Kit (V2 chemistry, 10X Genomics). For the subsequent three patient samples, libraries were prepared using the Single Cell 5’ Reagent Kit (Immunoprofiling Kit, 10X Genomics), with an additional step to enrich for V(D)J reads, following the manufacturer’s protocol.

#### GEM Generation and Barcoding

We loaded the chip to recover 10,000 cells (5,000 Tregs and 5,000 Tconvs) per sample.

#### Post-GEM Reverse Transcription and Library Preparation

These steps were performed according to the manufacturer’s instructions.

#### TCR Amplification

Single-cell TCR amplification and sequencing were performed after 5’GEX generation using the Single Cell V(D)J Kit, following the manufacturer’s protocol (10X Genomics). Briefly, V(D)J segments were enriched from amplified cDNA via two human TCR-targeted PCRs, followed by specific library construction.

#### Quality Control

Quality control of cDNA and V(D)J indexed libraries was conducted using a dsDNA High Sensitivity Assay Kit and the Bioanalyzer Agilent 2100 System. Libraries were then pooled equimolarly and sequenced accordingly.

#### cDNA Library Sequencing

cDNA libraries were sequenced using Illumina HiSeq and NovaSeq platforms in paired-end mode (26×98bp for transcriptome or gene expression), targeting at least 100,000 reads per cell.

#### V(D)J Library Sequencing

V(D)J libraries were sequenced on an Illumina HiSeq or MiSeq platform in paired end 150bp model, targeting at least 50,000 reads per cell.

### Pre-processing of scRNAseq

Raw base call (BCL) files produced by Illumina sequencer were demultiplexed and converted into Fastq files using cellranger mkfastq function from Cellranger version 2.1.1 with default parameters and bcl2fastq2 version 2.20. Generated Fastq files were processed using Cellranger version 3.0.2, that introduces an important improvement based on the EmptyDrops method^138^ to identify population of low RNA content cells. Cellranger count was run on each Fastq file based on 10xGenomics provided hg38/GRCh38 human reference genome (refdata-cellranger-GRCh38-1.2.0).

### scRNA-seq analysis

#### Filtering of data

downstream analysis was done using R package Seurat version 3.1.1-3.2.3 with R version 3.6.1-3.6.2^139,140^. After imported output files produced by cellranger into seurat pipeline, some filters were applied:

- Cells with fewer than 200 genes were removed to filter debris, death cells and other cells with few genes.
- A second filter has applied to remove other uninformative cells using a process based on percentage of mitochondrial genes and total counts of UMI by cell. In details, 1) we characterize the polymodal distribution of the cell counts by log2 percentage of mitochondrial genes and the polymodal distribution of cell counts by log2 total count of UMI by cell; 2) for each of these distributions, we algebraically determined the two maxima values of the polymodal distribution to identify the lowest minimum value between them. These minimum values will be use as cutoff to determine the uninformative cells. In the cases which no minimum could be identified for the distribution for mitochondrial genes, we used 10% as cutoff; 3) we filtered cells with percentages of mitochondrial genes higher than the cutoff defined by the polymodal distribution and with a total count of UMI by cell lower than the cutoff defined by the polymodal distribution of UMI by cell.

#### Elimination of Contaminant cells

data were normalized sample by sample using global-scaling normalization (using NormalizeData function with ‘‘LogNormalize’’ as normalization method) and the 2,000 most highly variable genes were identified for each sample (using FindVariableFeatures with ‘‘vst’’ as method, with low- and high-cutoffs for feature dispersions fixed at 0.5 and Inf and with low- and high-cutoffs for feature means fixed at –Inf and Inf).

To merge the different samples from 3 different tissues of 5 different patients, we used the integration process proposed by Seurat. This method consists in identifying “anchors” between different datasets (from different individuals, experimental conditions, …). These anchors make it possible to correspond cells from different samples, but cells sharing similar biological states, in order to harmonize them in a dataset. Integration process was performed with the identified anchors (using FindIntegrationAnchors followed by IntegrateData, both using the top 30 PCs). Integrated data was scaled (using ScaleData). No additional process has been done to correct batch effect. To reduce the technical noise, principal component analysis (PCA) was performed to work on the most contributing principal components (PCs). Graph-based clustering was done at different resolutions (using FindNeighbors on the first fifty PCs and FindClusters for the resolutions between 0 and 2 for each decimal) and visualized using Clustree version 0.2.2^141^. UMAP reduction was performed (using RunUMAP on the first fifty PCs) to visualize the data in UMAP projection. Clustering with resolution 0.1 was satisfying for the identification of contaminant cell based on absence of expression of T cell markers (*CD3E, CD3G, TRAC, TRBCA* and *TRBC2*) and expression of other immune population markers (CD79A for B cells, *CD14* for monocytes, *CD11C* for dendritic cells).

#### Integration

once identified contaminant cells were removed from data obtained after filters (Filtering of data step), and we repeated the analysis described above: normalization for each sample followed by highly variable gene identification. Then, samples were merged using integration process, data was scaled, graph-based clustering was done and UMAP reduction was performed (as described above).

#### Selection of resolution and clusters

Clusters obtained at resolution 1.1 were used for downstream analysis after merging clusters C2 with C19 (Tconv-N&Q).

Three clusters (PatA, PatB, and TRBV7) were excluded from the characterization due to low cell counts, TCR bias or because over 75% of the cells were contributed by only one patient.

#### Differential analysis

to characterize identities of cell clusters, differentially expressed genes among clusters were identified with MAST^142^ (using FindAllMarkers and returning only positive markers with a minimum log Fold-Change (logFC) of 0.2 and a minimum fraction of cells expressing the gene in either of the two groups (min.pct) fixed at 0.05).

Differential analyses summarized in **Table 2** were performed on all included genes using FindAllMarkers() function with following parameters: assay= “RNA”, min.pct= 0.05, logfc.threshold= 0, test.use= “MAST”, only.pos= FALSE, features= list_of_features, return.thresh= 1.

To characterize all the differential expressed genes by cluster, a paired-wise analysis was performed between each cluster and all the others (using FindMarkers with a minimum log Fold-Change (logFC) of 0.2 and a minimum fraction of cells expressing the gene in either of the two groups (min.pct) fixed at 0.05) and with adjusted p-value inferior or equal to 0.05.

To identify differentially expressed genes between Treg-Tfr and Tconv-N&Q, a pairwise analysis was performed using FindMarkers, with a minimum log fold-change (logFC) of 0.05 and a minimum fraction of cells expressing the gene in either group (min.pct) set at 0.05. All genes are displayed in the volcano plot, and genes with a logFC greater than or equal to 0.2 or less than or equal to −0.2, along with an adjusted p-value less than or equal to 0.05, are highlighted.

#### Signatures

in addition of differential analysis, signatures from public data were used (using AddModuleScore using 100 features as control and 24 bins (default parameters)).

#### Reassignment

for “state driven” clusters, which displayed a mixture of Treg and Tconv profiles, we used the transfer label function to differentiate cells between Tconv-like and Treg-like (using TransferData with the 30 most contributing PCs).

#### Ingenuity Pathway Analysis (IPA)

pathway analyses of differential expressed genes were uploaded on Ingenuity Pathway Analysis (Ingenuity® Systems, www.ingenuity.com) for the analysis of “Disease and bio-function”, “Canonical pathway”, “Causal network”, and “Upstream regulators”. Pathways were considered significantly when the overlap p ≤ 0.05.

#### Gene set Enrichment Analysis (GSEA)^143^

Gene set enrichment analyses were performed using fgsea() function of R package fastGSEA v1.26.0 (1) (modified parameters: with nPermSimple = 100,000) on log2 foldchange values (selected cluster versus all others), using FoldChange() function. All Gene set Enrichment analyses were performed on R environment v4.3.0 and using R package Seurat v4.3.0 for scRNA-seq data.

#### NeoAg and virus-signatures. Procedure adapted from^59^

Tumor single cells were ranked by module scores for neoantigen-specific and virus-specific gene signatures^59,60^. Cells in the 95th percentile, representing the top 5% with the highest gene signature expression, were classified as positive for tumor-specific or virus-specific signatures, respectively. Clonotypes corresponding to these cells were identified, and their tissue distribution was visualized using UMAP.

#### Velocity analysis

to investigate developmental dynamics in our data, Velocyto version 0.17.17^144^ with python version 3.6.2 was used to annotate reads between spliced, unspliced and ambiguous genes for each sample individually. Obtained loom files were processed using SeuratWrappers version 0.3.0, Seurat version 4.0.4, SeuratDisk version 0.0.0.9019 and velocyto.R version 0.6, with R version 4.1.1 and combined by tissue. For downstream analysis, RNA velocities were computed on python version 3.9.5-3.9.21 with scVelo stochastical and dynamical model (version 0.2.3-0.3.3^145^) using the 2,000 most variable genes from h5ad files generated from LNs and tumors separately. Both models were tested. Taking into account the confidence in the data vectors between the two models, we selected the best model which fit our data. RNA velocities analysis was projected into UMAP and diffusion map dimensional reduction plots using Destiny. In addition, Partition-based graph abstraction (PAGA) was used to optimize visualization of the directional orientation of RNA velocity in diffusion map projection.

#### R Shiny application (https://github.com/wilfridricher/scViz)

To facilitate the exploration of the single-cell data analysis presented in this paper, we developed an R Shiny application called scViz. This application visualizes clusters, cell density, and gene expression or signature scores in the desired two-dimensional space (such as UMAP and t-SNE). Additionally, gene or signature expression can be visualized using violin plots and heatmaps. This R Shiny application can also be applied to any Seurat object saved as .RData or .RDS files.

### scATAC-seq experiment

For the scATAC seq samples (Blood, LN, and Tumor from patients 6 and 7), nuclei were isolated and transposition and further steps were performed following manufacturer’s instructions (Chromium Single Cell ATAC Reagent Kits). Indexed ATAC-seq libraries were tested for quality (using a dsDNA High Sensitivity Assay Kit and the Bioanalyzer Agilent 2100 System), equimolarly pooled and sequenced with Illumina Nova-Seq, targeting at least 100,000 reads per nucleus.

#### GEM Generation and Barcoding

We loaded the chip to recover 10,000 nuclei (5,000 Tregs and 5,000 Tconvs) per sample.

### scATAC-seq analysis

Pre-processing of scATAC-seq: demultiplexing and conversion of raw base call (BCL) files into Fastq files were performed using bcl2fastq2 version 2.20 (with the following options: --no-lane-splitting --minimum-trimmed-read-length=8 --mask-short-adapter-reads=8 --ignore-missing-positions --ignore-missing-controls --ignore-missing-filter -- ignore-missing-bcls).

To align and quantify scATAC-seq reads, and to generate fragment files (containing all unique scATAC-seq fragments linked to the corresponding cellID) for the different samples, cellranger-atac count from Cellranger ATAC version 1.0 was used with the hg38/GRCh38 human reference genome provided by 10xGenomics (refdata-cellranger-atac-GRCh38-1.0.0).

#### Analysis of scATACseq

Downstream analyses were processed using R package ArchR version 0.9.5-1.0.1^45^ with R version 3.6.1. Annotation was done on hg38/GRCh38 human reference genome. Fragments files generated by Cellranger were imported in ArchR, low quality cells were discarded based on TSS enrichment (with a threshold fixed at 7) and fragments per cell (with a threshold fixed at 1,500). To resolve the technical differences observed in the lymph node sample from patient 2 compared to the other samples, down-sampling of the number of fragments was performed by taking only half of the fragments contained in this sample. Low quality cells of this sample were discarded as described above. Inferred doublets (using addDoubletScores function, a predictive approach proposed by ArchR which consists in the identification of nearest neighbor cells of in silico doublets from the data) were filtered after merging all samples.

#### Elimination of contaminant cells

iterative Latent Semantic Indexing (iLSI), a dimensionality reduction method, was applied on data composed of all good quality cells from the different samples (using addIterativeLSI based on the “TileMatrix” with 10 iterations on 50,000 cells and 5,000 variable features and passed the following parameters to the addClusters function: resolution = 0.1, sampleCells = 50,000, n.start = 10). Batch effect correction was performed using Harmony method^146^. Processed data were clustered using graph-based clustering implemented in Seurat at different resolutions (between 0 and 2 for each decimal) and UMAP reduction was computed to visualize the data. Contaminant cell subsets were defined using clustering resolution at 0.2, based on estimated gene expression (Gene-Scores) (approach based on the global contribution of chromatin accessibility within the entire gene) of T cell markers (CD3D) and expression of other immune population markers (CD14, CD8A, CLEC4C or MS4A1).

#### Characterization of scATAC cell populations

identified contaminant cells were removed from the data and analysis was repeated. iLSI was performed on good quality data (using addIterativeLSI based on the “TileMatrix” with the same parameters as before). Batch effect correction was applied using Harmony method. Processed data were clustered at different resolutions (between 0 and 2 for each decimal) and UMAP reduction was computed. Clusters obtained at resolution 0.9 were used for downstream analysis. To characterize identity of cell clusters, the identification marker features proposed by ArchR was performed on matrix Gene Score (using Wilcoxon test and considering only genes with a minimum logFC of 0.2 and a maximum false discovery rate (FDR) of 0.05).

To further characterize the difference between Treg-Tfr and Tconv-N&Q clusters, paired-wise identification marker features was performed on matrix Gene Score (using Wilcoxon test and considering differential markers only genes with a maximum false discovery rate (FDR) of 0.05 and a logFC superior or equal at 0.2.

##### Gene Score

A gene score is a metric used to predict gene expression based on the accessibility of regulatory elements near a gene, reflecting how likely the gene is actively expressed.

##### Positive Regulator

A positive regulator is a transcription factor identified by its enriched binding motif in accessible chromatin regions, whose expression correlates with the observed chromatin accessibility, indicating it likely drives gene regulation in those regions.

#### ScRNA-seq-scATAC-seq integration

to cross-compare cluster identities assigned using Gene Scores in scATAC-seq with identities characterized by gene expression in scRNA-seq, Seurat’s Transfer anchors process adapted by ArchR, which consist to identify “anchors” to correspond cells sharing similar biological states from different samples (from different technologies), was performed with addGeneIntegrationMatrix function using 15,000 reference cells from both data sources. Resulting cluster identities and gene expression of scRNA-seq cells were reported to the closest scATAC-seq cells.

#### Peak calling

peaks calling was performed using MACS2^147^ via the iterative overlap peak merging procedure proposed by ArchR (using addReproduciblePeakSet on cluster from resolution 0.9 with default parameters). Cisbp database was used to define and annotate peaks containing transcription factor (TF) motifs. Differential peaks among clusters were characterized using getMarkerFeatures function (using Wilcoxon test). From these differential peaks, first prediction transcription factor activity among clusters were performed (using peakAnnoEnrichment with a minimum logFC of 0.5 and a maximum FDR of 0.1) and visualized by heatmap (clustered using euclidean distance with Ward’s method). In a second time, ChromVAR^148^ deviation z-scores were calculated to predict TF enrichment for individual cells of the data. To predict the precise binding site of TF motif, ArchR’s getFootprints function was used.

#### Peak signatures

public peak list signatures were used as peak annotation (using addPeakAnnotations). To convert genomic coordinates of peaks from mm10 mouse genome or previous (hg18/NCBI36, hg19/GRCh37) human genome to hg38/GRCh38 human reference genome, we used the web version of LiftOver tool from UCSC^149^ (https://genome.ucsc.edu/cgi-bin/hgLiftOver). For peak signatures obtained on previous human genome reference, we have considered a ‘minimum ratio of base that must remap’ of 0.95, while a ‘minimum ratio of base that must remap’ of 0.2 was considered as sufficient for signatures characterized on mouse data. Then, we performed hypergeometric enrichment of these public signatures composed by LiftOver regions within the defined marker peaks in our data (using peakAnnoEnrichment, with a minimum logFC of 0.5 and a maximum FDR of 0.1) and results were visualized into an heatmap (using plotEnrichHeatmap). In addition, we calculated ChromVAR deviation z-scores.

#### Visualization

MAGIC^150^ (Markov Affinity-Based Graph Imputation of Cells) smoothing signal procedure, imputing weight to each cell, was applied on Gene Score, gene expression and deviation z-scores before visualization using UMAP reduction.

#### Co accessibility and Peak2Genes

co-accessibility between peaks of same gene among cells (co-accessibility) and co-accessibility between peaks and gene expression of same gene among cells (peak2genelinkage) were explored using ArchR’s functions (using addCoAccessibility following by getCoAccessibility and addPeak2GeneLinks following by getPeak2GeneLinks with dimension data obtained with Harmony). Briefly, co-accessibility highlights peaks which accessibility correlates across many single cells; while peak2genelinkage highlights peaks which accessibility correlates with gene expression.

#### Transcription Factor (TF) Footprinting

to visualize footprinting of specific TFs, we applied ArchR’s functions (getFootprints and plotFootprints with Substract normalization method) on two subgroups of clusters: “Clusters sharing Tissue Residency program” and “Other clusters”).

#### Regulatory network analysis

To construct regulatory network, scATAC-seq coupled to scRNA-seq were used. First, differential analysis was performed between naïve/memory, effector and tissue imprinted clusters to identify candidate target genes. In parallel, after selecting positive TFs based on Gene Integrated data, a binary matrix containing peaks scanned for these TF binding sites was extracted from data results obtained from ArchR (using getMatches). Then, the peaks contained in this matrix were linked to the union of candidate target genes using Peak2Genes output (using getPeak2GeneLinks function with the following parameters: corCutOff = 0.5, resolution = 10,000, returnLoops = FALSE). To further define the regulatory network, Pearson correlations between candidate target genes and TFs were performed on scRNA-seq data. Low and/or non-significant correlations (|Pearson correlation| > 0.1 and p-value < 0.001) and “TF – candidate target gene” pair without peak detection were removed. Finally, the regulatory network was designed from these results using Gephy.

#### Trajectory analysis

taken into consideration results from scRNA-seq trajectories and velocity analysis, supervised trajectory analysis in all tissues and per tissue (blood only, lymph node only and tumor only) were performed using ArchR’s procedure. ArchR’s supervised trajectory analysis determines a coordinate and a pseudotime value for each cell of the designed trajectory, based on distance between each cell and the next cluster in the trajectory. Once done, gene scores, gene expression, deviation z-scores and peak accessibility were visualized by heatmap and UMAP. Integrative analysis between correlated gene score and TF motif accessibility or between gene expression and TF motif accessibility were performed (using same parameters than ArchR manual) to produce paired heatmaps for the corresponding features.

#### Overall results

Each nucleus yielded on average of 11.5 X 10^3^ unique fragments mapping to the nuclear genome, and a fraction of 0.6 reads in peak regions. Around 7,317 peaks/cell and 8,347 predicted genes/cells (gene expression estimated on the accessibility of regulatory elements in the vicinity of the gene, gene score) were obtained, compared to the 1,260 genes/cell obtained by scRNA-seq.

### Flow cytometry analysis of follicular subsets

#### Sample preparation

Single-cell suspensions were prepared as previously specified and frozen in 90% bovine serum and 10% DMSO, following standard protocols. Prior to the experiment, they were thawed in CO2-independent medium (GIBCO) containing 0.4% human BSA at 37 °C and quickly washed twice with medium to minimize cell death.

#### FACS staining

Cell suspensions were washed in PBS and incubated with LIVE/DEAD Fixable Cell Dead Stain (eBioscience) during 15 min at RT. Next, cells were washed, resuspended to a concentration of 30.10^6^ Cells/mL, and incubated with fluorochrome labeled Abs for 30 min at 4°C. Finally, the cells were washed and resuspended in MACS buffer to a final concentration of 30.10^6^ Cells/mL for sorting or kept for intracellular staining.

#### Intranuclear staining

Cells were fixed, permeabilized and stained with Foxp3/Transcription Factor Staining Buffers (eBioscience) following eBioscience One-step protocol manufacturer’s instructions. In the case of unlabeled Abs, we included an additional step where we stained the samples for 30 minutes with anti-specie specific Abs, that recognized the primary antibodies and were conjugated with fluorochromes. Data was acquired with a BD LSR-Fortessa flow cytometer and exported into FlowJo software (version 10.0.8) and GraphPad Prism v10.0.2.

### FACS-sorting and bulk RNA/TCR-seq experiment

#### Sample Preparation

We sorted Treg-Tfr, Treg-Trm, Treg-N, Tconv-Tfh, Tconv-CXCL13, and Tconv-N populations from frozen samples of 5 patients, collected from two different tissues: lymph nodes (LNs) and tumors. Cells were initially frozen in 90% bovine serum and 10% DMSO, following standard protocols. Prior to the experiment, they were thawed in CO2-independent medium (GIBCO) containing 0.4% human BSA at 37 °C and quickly washed twice with medium to minimize cell death.

#### FACS sorting

To sort the specified populations, we employed the previously described gating strategy and procedure (FACS characterization of follicular CD4+ T cells, without intracellular staining), further refined by the addition of ICOS anti-human antibody to better discriminate effector and Trm Tregs from Treg-Tfr. This enhancement allowed us to better distinguish Tconv-CXCL13 and Treg-Trm clusters from Treg-Tfr. During the experiment, cells were kept in MACS buffer at 4°C and directly collected into 1,5mL Eppendorf containing TLS buffer containing 1% beta-mercaptoethanol (Qiagen). Of note, some cells were sorted in MACS buffer to corroborate FOXP3 expression (following FACS characterization of FTA-CD4+ T cells procedure) in sorted Treg populations. Cell-sorting was performed using ARIA Fusion cell sorter (BD). Equal numbers of cells were sorted for each population per tissue/patient, and the samples were stored at −80°C until RNA purification.“

#### RNA purification, library construction, and sequencing

RNA was isolated using the single cell RNA purification kit (Norgen). RNA quality was assessed using the picoRNA kit (Bioanalyzer) and concentration measured with the Qubit RNA HS Assay Kit (Thermo Fisher Scientific). Then, total RNA was split in equal parts for RNA-seq and TCR-seq retro transcription, amplification, and library construction.

#### RNA libraries

RNA was retrotranscribed and amplified using the SMART-Seq v4 PLUS Kit (Takara). cDNA libraries were constructed following the manufacturer’s recommendations. The quality of the cDNA and final libraries was controlled and measured using HS-DNA kit (Bioanalyzer) and the Qubit RNA HS Assay Kit (Thermo Fisher Scientific), respectively. Finally, bulk RNA-seq samples were sequenced on a Novaseq (Illumina) with 2 × 100 bp read length aiming for 20 million reads per RNAseq sample

#### TCR libraries

RNA was retrotranscribed and amplified using the Human TCR RNA multiplex kit (Mi Laboratories). Importantly, TCR RNA multiplex kit includes UMI labeling along with 1st strand cDNA synthesis to allow for advanced PCR and sequencing error correction, elimination of PCR biases, exact quantification of template cDNA molecules, and accurate normalization of samples for comparison of repertoire diversity metrics. cDNA libraries enriched in TCR transcripts were constructed following the manufacturer’s recommendations. The quality of the cDNA and final libraries was controlled and measured using HS-DNA kit (Bioanalyzer) and the Qubit RNA HS Assay Kit (Thermo Fisher Scientific), respectively. Finally, samples were sequenced on a Novaseq (Illumina) with 150 bp for TCR libraries, aiming for at least 100 paired-end reads per cell.

### Bulk RNA-seq analysis

The count matrix was analyzed using the DESeq2 pipeline (R package version 1.38.3). MicroRNAs (miRNAs), mitochondrial, and ribosomal genes were excluded from the dataset. Genes with more than 50 counts in at least three samples were retained for downstream analysis. Normalization was performed using the median of ratios method, as described by Anders and Huber (2010)^151^. Batch effects were corrected using the removeBatchEffect() function from the Limma package (version 3.54.2). Hierarchical clustering was conducted with the “ward.D2” method, considering the 3,000 most variable genes. For differential expression analysis, we used the DESeq() function and performed hypothesis testing using a Wald test^152^. We then used the FDR (Benjamini-Hochberg) method for the multiple testing correction. The rlog transformation was used for visualization. The heatmap represents z-scores of normalized gene expression.

#### Gene set Enrichment Analysis (GSEA)^143^

Gene set enrichment analyses were performed using fgsea() function of R package fastGSEA v1.26.0 (1) (modified parameters : with nPermSimple = 100,000) on log2 foldchange values (selected cluster versus all others). All Gene set Enrichment analyses were performed on R environment v4.3.0.

### scTCRseq analysis

#### Pre-processing of scTCR-seq

Fastq files were produced from raw base call (BCL) files using cellranger mkfastq function from Cellranger version 2.1.1 with default parameters and bcl2fastq2 version 2.20. To produce TRA and TRB V(D)J sequence assembly, each generated Fastq files was processed using Cellranger vdj function from Cellranger version 3.1.0 based on 10xGenomics provided hg38/GRCh38 human reference genome (refdata-cellranger-vdj-GRCh38-alts-ensembl-2.0.0).

#### Analysis of scTCR-seq

processed files by Cellranger were imported in R and clones without full-length and non-productive contigs (which compose the clonotypes of Cellranger) were discarded for downstream analysis. To improve the definition of clonotypes and considering that TRA and TRB chains are defined by combining CDR3 and V and C subunits, we considered for our analysis only those cells presenting only 1 TRA chain and 1 TRB chain to define clones. To estimate clonal diversity, we used Gini-TCR Skewing Index^153^ and Morisita-Horn index.

#### Morisita-Horn Index (MHI) coupled to permutation test

To estimate TCR sharing between two different cell groups, we performed Morisita-Horn Index using the following formula:

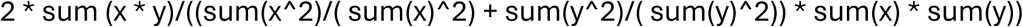

Where x is the number of occurrences of each clonotype observed in the population A, and y is the number of occurrences of each of the same clonotypes observed in the population B.

To determine the probability of an observed Morisita-Horn iIndex (MHI) being different from random, we used a permutation test. We randomized the TCRs between both cell types and recalculated the MHI post-randomization. This process generated empirical MHI distributions through 1000 permutations, which served as the null distribution of MHI. We then tested if the observed MHI was significantly different from the randomized scenario (MHI null distribution) using a two-tailed empirical p-value approach.

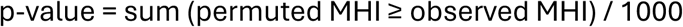

This approach allows us to assess whether the observed overlap in TCR repertoires between two cell types is significantly different from what would be expected by chance, providing evidence for or against shared clonal relationships between the populations.

Considering that most of the TCRs in our dataset were singlets or low-expanded clones, and that the studied sharing phenomena — such as Treg/Tconv repertoire overlap or conversion, Lymph Node/Tumor overlap or migration, and Treg/Treg or Tconv/Tconv transitions — were relatively rare events, we restricted our analysis to focus on overall sharing patterns, as follow. To reduce noise, we excluded blood-derived cells from all comparisons. Specifically:

- For the Treg/Tconv TCR sharing, we identified clonotypes at the intersection between Treg and Tconv cells and used only these cells (as 100% of the dataset) for further calculations. Among all TCRs from “pure” Treg clusters Treg_N&Q, Treg_Trm and Treg_Tfr (1,774 clonotypes) and all TCRs from “pure” Tconv clusters Tconv_N&Q, Tconv_Tfh and Tconv_CXCL13 (2,105 clonotypes), we found only 36 shared clonotypes. For subsequent using MHI and permutation (p-value) calculation as previously described, we restricted our focus to the cells belonging to these shared clonotypes (57 and 52 cells for Treg and Tconv clusters respectively).
- For the LN/Tumor TCR sharing, we identified clonotypes at the intersection between LNs and tumor cells and used only these cells (as 100% of the dataset) for further calculations. Among all TCRs from LNs (2,574 clonotypes) and all TCRs from tumors (1,363 clonotypes) for clusters previously cited (Treg_N&Q, Treg_Trm, Treg_Tfr, Tconv_N&Q, Tconv_Tfh and Tconv_CXCL13), we found only 94 shared clonotypes. For subsequent using MHI and permutation (p-value) calculation as previously described, we restricted our focus to the cells belonging to these shared clonotypes (141 and 187 cells for LNs and tumors respectively).

This approach allowed us to increase the statistical power for detecting these infrequent events.

#### TCR databases

The defined clonotypes were then compared to different databases (McPAS-TCR and VDJdb, August 2024) by accepting up to 1 amino acid mismatch on the TRB. The TRA was not considered. Cells with clonotype references in the databases were represented in UMAP according to three categories based on their associated Ag: Pathogenic or Autoimmune. R, Excel and Prism 10 software were used for calculation, statistical analysis and visualizations.

### Bulk TCR-seq analysis

#### Bulk TCR-seq analysis

Fastq files were processed using MixCR v4.5.0 for milab-human-rna-tcr-umi-multiplex experiments preset on python 3.9. Clonotypes were defined using CDR3/V subunit. We count the number of UMI and different clonotypes obtained for each sample.

To estimate clonal similarity between groups of samples, we performed Morisita-Horn index coupled to a permutation test to identify the most significant results (detailed description in scTCR-seq part).

#### Morisita-Horn index (MHI) coupled to permutation test

We follow the same procedure than for the single-cell TCR-seq. The clonotypes and UMIs considered for each comparisons considering all patients together were:

| A/B | Clonotypes A | Clonotypes B | Shared Clonotypes (SC) | UMIs comprised on SC |
| --- | --- | --- | --- | --- |
| **Treg/Tconv** | 27,148 | 17,202 | 1,737 | 22,796<br>(Treg=8,315/Tconv=14,481) |
| **LNs/Tumor** | 33,930 | 10,206 | 1,523 | 18,903<br>(Tum=9,302/LN=9,601) |

### In Vitro Tumor Antigen Specificity Assay

Single-cell suspensions (1 × 10⁶ total cells) were prepared from tumor-invaded lymph nodes (LNs), which naturally contain both antigen-presenting cells (APCs) presenting tumor-derived antigens and tumor-infiltrating T cell populations. Cells were cultured in X-VIVO medium (Lonza) supplemented with 10% human serum for 16 hours at 37°C in a humidified incubator with 5% CO₂. Cultures were established in duplicate; one condition included an anti-human HLA-DR blocking antibody (Miltenyi Biotec, 100 µg/mL) to inhibit MHC class II-mediated antigen presentation.

Following incubation, which allowed antigen presentation and subsequent T cell activation and consequent 4-1BB (CD137) upregulation, cells were harvested, washed with PBS, and stained for flow cytometric analysis. Staining included a viability dye (LIVE/DEAD Fixable Aqua, Thermo Fisher) and fluorochrome-conjugated antibodies specific for T subpopulation markers and 4-1BB.

Flow cytometry was performed on a BD LSR-Fortessa, and data were analyzed using FlowJo v10.9.0. Statistical analysis was conducted using GraphPad Prism v10.0.2.

### Treg Identity definition using transfer label function

Treg-Tfr cells were classified as Tregs or Tconvs using Seurat’s transfer label function. Previously defined Treg and Tconv clusters (excluding state-driven clusters) were used as reference populations. Prediction scores were calculated following Seurat’s pipeline instructions for each cell in the Treg-Tfr cluster to assign cell identity.

### In Vitro Treg Fragility Assay

Treg and Tfr cell subsets were isolated from healthy donor peripheral blood mononuclear cells (HD-PBMCs) following CD25+ cell enrichment using a magnetic selection kit (Miltenyi Biotec) following manufacture instructions. Enriched cells were subsequently staining with viability markers and CD4, CD45RA, CD25, CD127 and CXCR5 antibodies and then sorted on a BD FACS ARIA Fusion flow cytometer into the following populations:

- **Treg-Tfr cells**: viable CD4⁺CD25^high^CD127^low^CXCR5⁺
- **Effector memory Tregs**: viable CD4⁺CD25^high^CD127^low^CXCR5⁻CD45RA⁻
- **Naïve Tregs**: viable CD4⁺CD25^high^CD127^low^CXCR5⁻CD45RA⁺

Following sorting, cells were counted, and 5 × 10⁴ cells per condition were plated into flat-bottom 96-well plates pre-coated with anti-CD3 (5 µg/mL in PBS, incubated for 2 hours at 37°C, followed by two PBS washes). Cells were then cultured for 3 days in X-VIVO medium (Lonza), supplemented with 10% human serum, non-essential amino acids (Gibco 1X), Penicillin-Streptomycin (Gibco 50IU/mL) and soluble anti-CD28 (1 µg/mL), under the following stimulation conditions:

- **Treg-polarizing cocktail**: IL-2 (300 IU/mL) and TGF-β (10 ng/mL)
- **Tconv-polarizing cocktail**: IL-6 (50ng/mL) and IL-1β (20ng/mL)
- **Unstimulated control**: (300 IU/mL)

After 3 days of culture, cells were washed to remove cytokines and anti-CD28, and incubated overnight in fresh medium. The following day, cells were stained with a viability dye, fixed, and intracellularly stained for FOXP3. Flow cytometry was performed on a BD

LSR-Fortessa cytometer, and data were analyzed using FlowJo v10.9.0. Statistical analyses were conducted using GraphPad Prism v10.0.2.

The experiment was independently replicated using samples from three additional donors in Dr. Graça’s laboratory (Faculdade de Medicina, Universidade de Lisboa, Lisbon, Portugal), with consistent results observed in two of the analyzed subjects.

### Multiplex Immunohistochemistry

#### Tissue preparation

Tumor and lymph node tissue samples were obtained from Institute Mutualist Mountsouris (IMM) and prepared by the platform of Experimental pathology of Institut Curie.

#### Multiplex IHC stating

Multiplex Immunohistochemistry (IHC) was performed according to the protocol developed by (Remark R and al. 2016^154^), with some adjustments: Formalin-fixed, paraffin-embedded (FFPE) tumor and lymph node tissue samples were sectioned at 3 µm thickness and mounted on charged glass slides. Tissue sections were baked for 1h at 60°C, then deparaffinized and rehydrated in successive baths of xylene and descending gradients of ethanol (100%,90%,70%,50%, diH2O). Heat-induced epitope retrieval was performed with pH6.1 (Agilent Dako, S169984-2) or pH9 (Agilent Dako, S236784-2) buffer, in a water bath at 95°C for 30 min for the first staining (otherwise 15 min).

Tissues were stained manually using the following protocol:

Peroxidase Blocking: Tissue slides were incubated in peroxidase blocking solution (Agilent Dako, S202386-2) for 10 minutes to quench endogenous peroxidase activity.

Fab Fragment Blocking (If required): If a primary antibody was of the same specie as one previously used in the staining protocol, Fab fragment blocking was applied for 20 minutes to prevent cross-reactivity.

Protein Blocking: After Fab blocking, a protein block (Agilent Dako, S202386-2) was applied for 10 minutes to block non-specific protein binding.

Primary Antibody Incubation: The slides were incubated with primary antibodies for 1–2 hours at room temperature. The following primary antibodies were used in this study:

|  |  |  |  |  |  |  |
| --- | --- | --- | --- | --- | --- | --- |
| 1 | FOXP3 | pH 9, 30 min, water bath | 1/50, 2h @RT | Mouse, 30 min @RT | 45min | clone 236A/E7, Abcam, cat# ab20034 |
| 2 | PD1 | pH 6, 15 min, water bath | 1/150, 2h @RT | Rabbit, 30 min @RT | 30 min | PD1 (polyclonal, Sigma Aldrich), cat# HPA035981-100UL |
| 4 | CD8 | pH 9, 15 min, water bath | 1/50, 2h @RT | Mouse, 30 min @RT | 30 min | Clone C8/144b, Agilent Dako, cat# M710301-2 |
| 5 | CXCL13 | pH 6, 15 min, water bath | 1/500 1h at RT | Goat, 30min @ RT | 30 min | Polyclonal, BioTechne (R&D), cat #AF801 |
| 7 | CD4 | pH 9, 15 min, water bath | 1/75, 2h @RT | Mouse, 30 min @RT | 30 min | Clone N1UG0, eBioscience, cat#14-2444-82 |
| 8 | CD200 | pH 9, 15 min, water bath | 1/75, 2h @RT | Rabbit, 30 min @RT | 30 min | Clone [EPR22412-229], Abcam, Cat# ab254193 |

Secondary Antibody Detection: After primary antibody incubation, slides were washed with TBS-Tween (TBST) and incubated with horseradish peroxidase (HRP)-conjugated secondary antibodies: Anti-rabbit: Agilent Dako, K400311-2; Anti-mouse: Agilent Dako, K400111-2; Anti-goat: BioTechne, VC004-050.

Chromogenic Detection: The HRP activity was visualized using a chromogenic substrate, 3-amino-9-ethylcarbazole (AEC) (Abcam, ab64252), which produces a red precipitate at the site of antibody binding.

Counterstaining and Mounting: Tissue sections were counterstained with hematoxylin (Agilent Dako, S330930-2) for nuclear staining, then mounted using a glycerol-based aqueous mounting medium (Agilent Dako, C056330-2) to preserve the tissue and staining for imaging.

Image Acquisition and Scanning: Stained slides were scanned using the NanoZoomer S360 digital slide scanner (Hamamatsu, C13220-01) at 20x magnification to obtain high-resolution images of the tissue sections.

Stain Removal and Next Staining Cycle: After scanning, the slides underwent ethanol baths to remove the staining, and the staining cycle was repeated, starting with heat-induced epitope retrieval in Tris-EDTA buffer (pH 9.0) for 20 minutes. This allowed for sequential staining of additional markers in the multiplex panel.

Overlay Steps: All stains from the same sample were deconvolved into a pseudo-fluorescence signal using the HALO Indicalab Deconvolution v1.1.8 module. Subsequently, the images were registered with the HALO Registration Module and fused into a single overlay in .tif format. Finally, the overlays were cropped and exported as .ome.tiff files for analysis using QuPath software.

Image Analysis Using QuPath Following multiplex staining, tissue images were analyzed using QuPath software (v0.4.3). The following steps were performed:

1. Image Import and Pre-processing: Scanned images were imported into QuPath, and color deconvolution was used to separate chromogenic signals for different markers.
2. Tissue annotation: Germinal Center in LNs and TLS (tumors) were selected according to CD200 and CD4 staining and were annotated as PD1high or PD1low according to PD1 staining. Paracortical zones or non-TLS zones, both enriched in CD4+ T cells, were selected as controls.
3. Cell Detection (using Stardist): Cells were detected using the hematoxylin-stained nuclei. A size filter was applied to exclude non-cellular objects.
4. Classifier: Single-measurement and composite classifier were trained using cell-mean staining for all the channels, except for Foxp3 were we used nucleus classifier.
5. Phenotyping: Detected cells were classified based on marker expression from the chromogenic staining. Composite images were used to classify specific cell types (e.g., CD4+FOXP3+CD200+PD1+ Treg-Tfr cells and CD4+FOXP3-CD200+PD1+ Follicular Tconv cells).
6. Spatial Analysis Quantification: Cell density in manually selected regions were quantified using QuPath’s spatial analysis tools.
7. Data Export and Statistical Analysis: Quantified data were exported for statistical analysis using GraphPad Prism, with results presented as % of cells.

### Spatial Transcriptomics analysis

We downloaded the dataset corresponding to sample ID V1_Breast_Cancer_Block_A_Section_1 from the 10X Genomics website. The sample is annotated as “Breast Cancer: Ductal Carcinoma in Situ, Lobular Carcinoma in Situ, Invasive Carcinoma” and was published on 2020-06-23. The dataset was normalized using the SCTransform() function in Seurat. To account for dropout effects, we applied the RunALRA() function from Seurat. Gene signature scores were computed using the AddModuleScore() function.

#### Curated Signatures for Spatial Transcriptomics

Gene expression signatures were curated from our single-cell RNA sequencing (scRNA-seq) data to identify differentially expressed genes (DEGs) specific to the indicated T cell populations and broader subsets (e.g., follicular or tissue-resident T cells). To ensure specificity, we selected genes that were uniquely or predominantly expressed within each population or subset in our scRNA-seq dataset. These candidate marker genes were then cross-referenced against additional spatial transcriptomics datasets and comprehensive tumor microenvironment (TME) datasets to exclude genes with significant expression in non-T cell compartments such as fibroblasts, tumor cells, or other stromal components. Only genes with T cell–specific expression profiles were retained for downstream spatial analyses. The final set of selected genes is indicated in the corresponding figure panels.

Dataset: https://www.10xgenomics.com/datasets/human-breast-cancer-block-a-section-1-1-standard-1-1-0

### Use of AI Tools

Large language models, including ChatGPT (OpenAI, GPT-4) and Claude (Anthropic), were used to assist in refining the wording, grammar, and clarity of sections of the manuscript, including the introduction and methods. All scientific content, analysis, and conclusions were conceived, conducted, and validated by the authors.

## SUPPLEMENTARY TABLES

**TABLE 1. Patients’ characteristics.**

**TABLE 2. DEGs of scRNAseq clusters.**

**TABLE 3. scRNAseq associated signatures.**

**TABLE 4. DEGs of bulk RNAseq subpopulations.**

**TABLE 5. DEGS of scATACseq clusters.**

**TABLE 6. DE peaks of scATACseq clusters.**

**TABLE 7. TFs and related TFBM.**

**TABLE 8. TF-regulators and target genes.**

**TABLE 9. scTCR-seq information.**

**TABLE 10. Bulk TCR-seq information.**

**TABLE 11. scATACseq signatures.**

## ACKNOWLEDGEMENTS

We thank Sylvain Baulande, Virginie Reynal, Benoit Albaud, Laura Baudrin, and Patricia Legoix at the Curiecoretech Next Generation Sequencing (ICGEX) plattform at Institut Curie. High-throughput sequencing was performed by the ICGex NGS platform of the Institut Curie supported by the grants ANR-10-EQPX-03 (Equipex) and ANR-10-INBS-09-08 (France Génomique Consortium) from the Agence Nationale de la Recherche (“Investissements d’Avenir” program), by the ITMO-Cancer Aviesan (Plan Cancer III) and by the SiRIC-Curie program (SiRIC Grant INCa-DGOS-465 and INCa-DGOSInserm_12554). Data management, quality control and primary analysis were performed by the Bioinformatics platform of the Institut Curie. We acknowledge Coralie Guerin, Anna Chipont, Annick Viguier, and Lea Guyonnetin at Curiecoretech Cytometry platform at Institut Curie. We thank Nathalie Amzallag, Sophie Viel, and Jeremie Goldstein for technical assistance. We acknowledge Dr. Leticia Niborski for the help with scATAC-seq experiments. We thank Sarah Lagah and Simon Lefranc for the help with the clinical samples. This work was supported by ICGex NGS platform (Institut Curie) and Egle therapeutics.

## AUTHOR CONTRIBUTIONS

J.T.B., W.R., J.W. and E.P conceptualization.

J.T.B., W.R., E.B., Y.M., S.L., M.P., J.D., J.W., and E.P. designed experiments.

L.M., S.G.A, C.S., N.G., clinical samples collection.

J.T.B, W.R., E.B., Y.M., S.L., M.P., R.N.R., F.R., and J.D. performed the experiments.

J.T.B, W.R., Y.M., S.L., M.P., J.W., and E.P analyzed scRNA-seq data.

J.T.B, W.R., Y.M., S.L., M.P., J.W., and E.P analyzed scATAC-seq data.

J.T.B., W.R., S.L., J.W., and E.P wrote the manuscript.

J.W., and E.P supervision.

S.K., L.G, and O.L. meaningful discussions and correction of the manuscript.

S.B., M.B., S.L. provided NGS support.

J.T.B., W.R., C.S., J.W., and E.P. funding acquisition.

## COMPETING INTEREST

E.P. is co-founder of Egle-Tx.

S.L. and M.P. are employees from Egle-Tx.

J.T.B. and J.W. were consultants for Egle-Tx.

## DATA AND MATERIAL AVAILABILITY

Data and material availability information will be provided in accordance with journal requirements before publication.

### Data availability

Tosello, Richer et al. dataset (NSCLC) has been deposited in EGA, with accession code EGAS50000000293 for scRNAseq and EGAS50000000294 for scATAC-seq. The sequence data are generated from patient samples and therefore are available under restricted access. Data access can be granted via the EGA with completion of an institute data transfer agreement, and data will be available for one year once access has been granted. In-house bulk-RNAseq has been deposited in GEO database with the GEO accession number: (in process).

---

**Supplementary Fig. S1.**
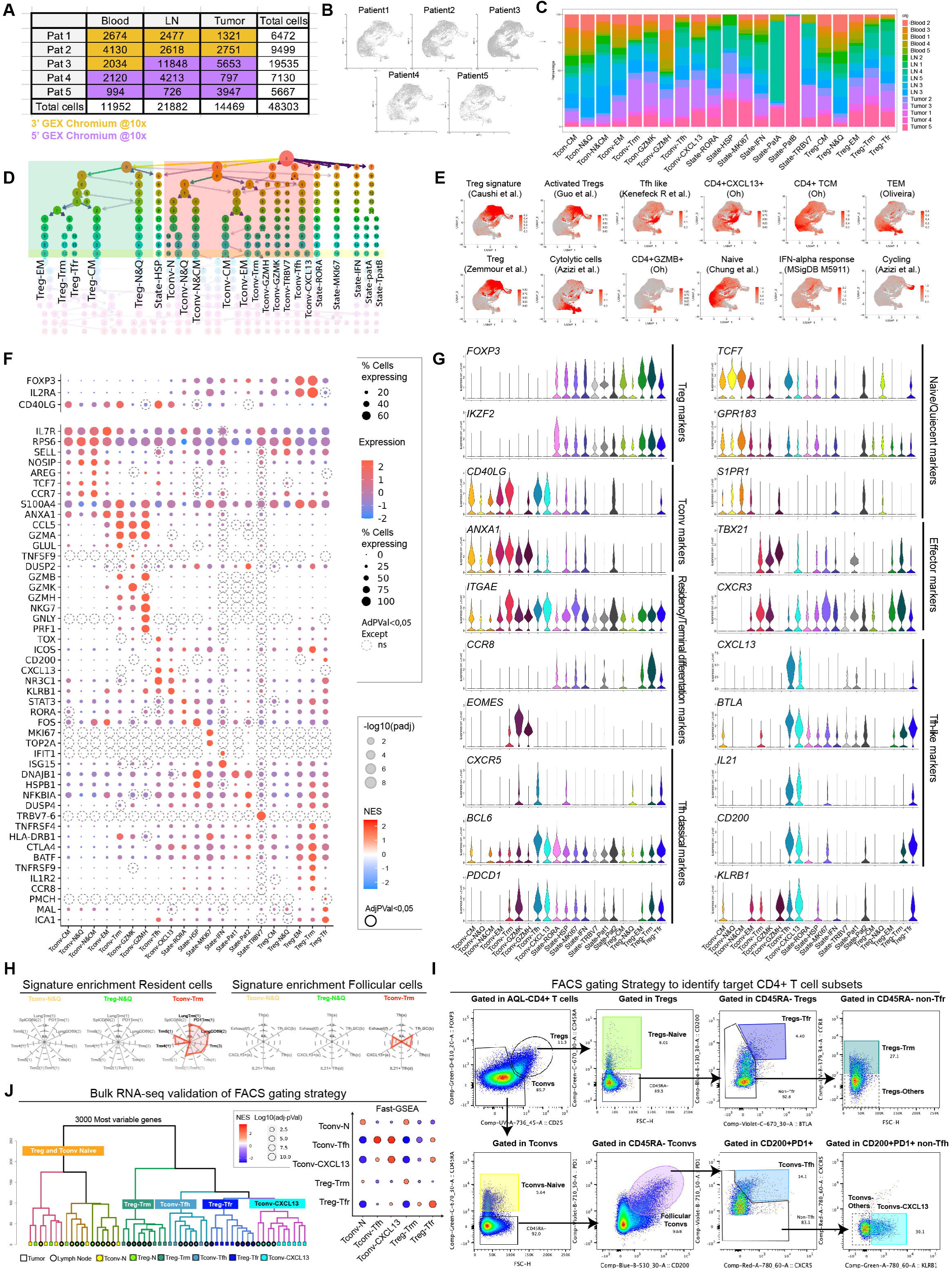
scRNA-seq analysis of CD4⁺ T cell populations across tissues. **(A)** Table displaying the number of cells per sample, color-coded by the 10X technology applied. **(B)** UMAP projection of total cells from all tissues, separated by patient. **(C)** Heatmap showing the relative proportions of cluster sizes across tissues: blood (B), lymph node (LN), and tumor (T). **(D)** Clustering tree plot tracking clusters across multiple scRNA-seq resolutions. **(E)** UMAP plots displaying gene signature scores (gray: low expression; red: high expression) for 12 CD4⁺ T cell population signatures (see **Table 3** for gene signature list and references)**. (F)** Dot plots illustrating the average expression (color intensity) and percentage (circle size) of cells expressing selected genes defining the 21 CD4⁺ T cell clusters. Genes with differential expression (p < 0.05) are represented, while non-significant ones (p > 0.05) are outlined with a circle. **(G)** Violin plots showing the expression distribution of selected genes across scRNA-seq-defined clusters. **(H)** Spider plots displaying NES values of Tconv-N&Q, Treg-N&Q and Tconv-Trm for resident and follicular cells signatures. Signatures with p < 0.05 are outlined in black, while non-significant ones are shown in gray. Signature references as in Fig. 1G. **(I)** Dot plots depicting different antibody staining and the sequential gating strategy used to identify follicular T cell subsets and control populations, including Tconv-CXCL13, Tconv-Tfh, Tconv-N, Tconv-Others, Treg-Trm, Treg-Tfr, Treg-N, and Treg-Others. **(J) Left:** Hierarchical clustering of the 3000 most variable genes (MVGs) distinguishing sorted populations. LNs are represented as circles, and tumors as squares. **Right.** Gene set enrichment dot plots showing Normalized Enrichment Score (NES) and p-values for each sorted population. The analysis compares bulk RNA-seq data with scRNA-seq-derived signatures for each cluster.

**Supplementary Fig. S2:**
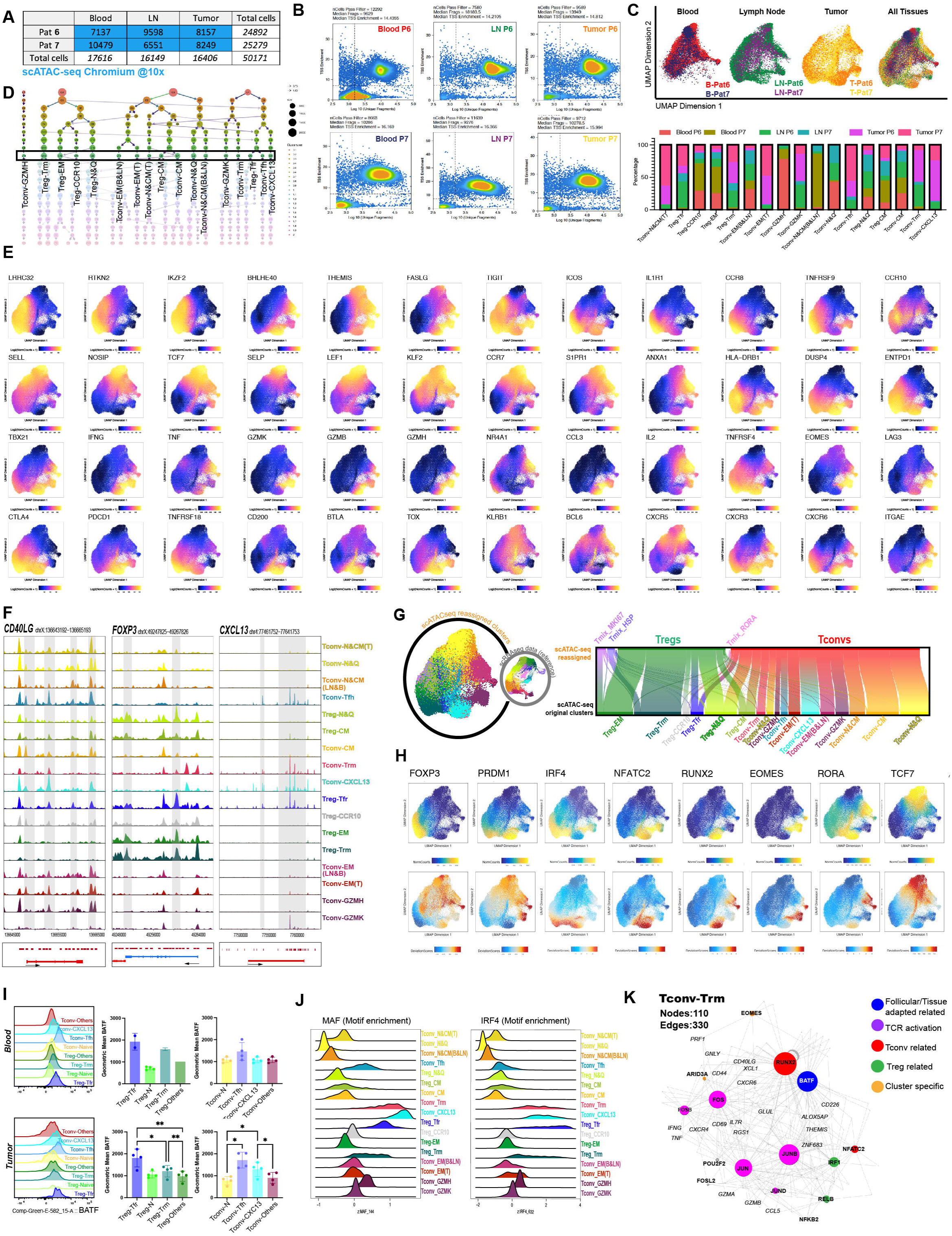
scATAC-seq analysis of CD4⁺ T cell populations across tissues. **(A)** Sample overview: Table summarizing the number of cells per sample analyzed using 10x Genomics scATAC-seq technology. **(B)** Quality control metrics: Dot plots showing per-cell quality control measures, including TSS enrichment and unique fragment distribution, for each sample. **(C)** Cell distribution across patients and tissues: (**Top**) UMAP projection of nuclei, colored by sample and split by tissue type. (**Bottom**) Bar plots quantifying the percentage of cells contributed by each sample. **(D)** Cluster hierarchy analysis: Clustering tree plot displaying the relationship of clusters across multiple scATAC-seq resolutions. **(E)** Gene score visualization: UMAP plots depicting gene score levels for 48 characteristic CD4⁺ T cell genes. Color scale represents expression levels (blue = low, yellow = high). **(F)** Chromatin accessibility at key loci: Genome track visualization of normalized peak accessibility at selected gene loci for each CD4⁺ T cell subset. Gene bodies are displayed at the bottom, with highlighted peaks in grey. **(G)** Integration of scATAC-seq and scRNA-seq data: **(Left)** UMAP plots comparing scRNA-seq cluster labeling (small circles) with scATAC-seq nuclei reassigned and colored based on RNA-seq-defined clusters (large circles). **(Right)** Alluvial plot illustrating the correspondence between scATAC-seq reassigned nuclei (categorized as pure Tconv, pure Treg, or state-driven clusters) and their original scATAC-seq clusters. **(H)** Transcription factor analysis: UMAP plots showing (top) gene integrated expression and (bottom) TF motif enrichment for eight selected TFs. **(I)** FACS analysis. **Left:** Histograms of BATF expression in gated populations from Blood and Tumor. **Right:** Bar plots quantifying BATF geometric mean fluorescence intensity in subsets from a representative NSCLC LN sample (n = 4 patients). Statistical test: One-way ANOVA (paired) with Tukey’s multiple comparisons post hoc test (* = p ≤ 0.0332; ** = p ≤ 0.0021; *** = p ≤ 0.0002; **** = p ≤ 0.0001). **(J)** MAF and IRF4 motif enrichment: Visualization of motif enrichment deviation scores for *MAF* and *IRF4* across clusters. **(K)** TF-target gene regulatory network: Network representation of positive TF regulators (from Fig. 2D) in **Tconv-Trm cluster**. Node size corresponds to out-degree, and node color indicates association with TCR activation, tissue-imprinted programs, Tconv/Treg subset identity, or cluster specificity.

**Supplementary Fig. S3:**
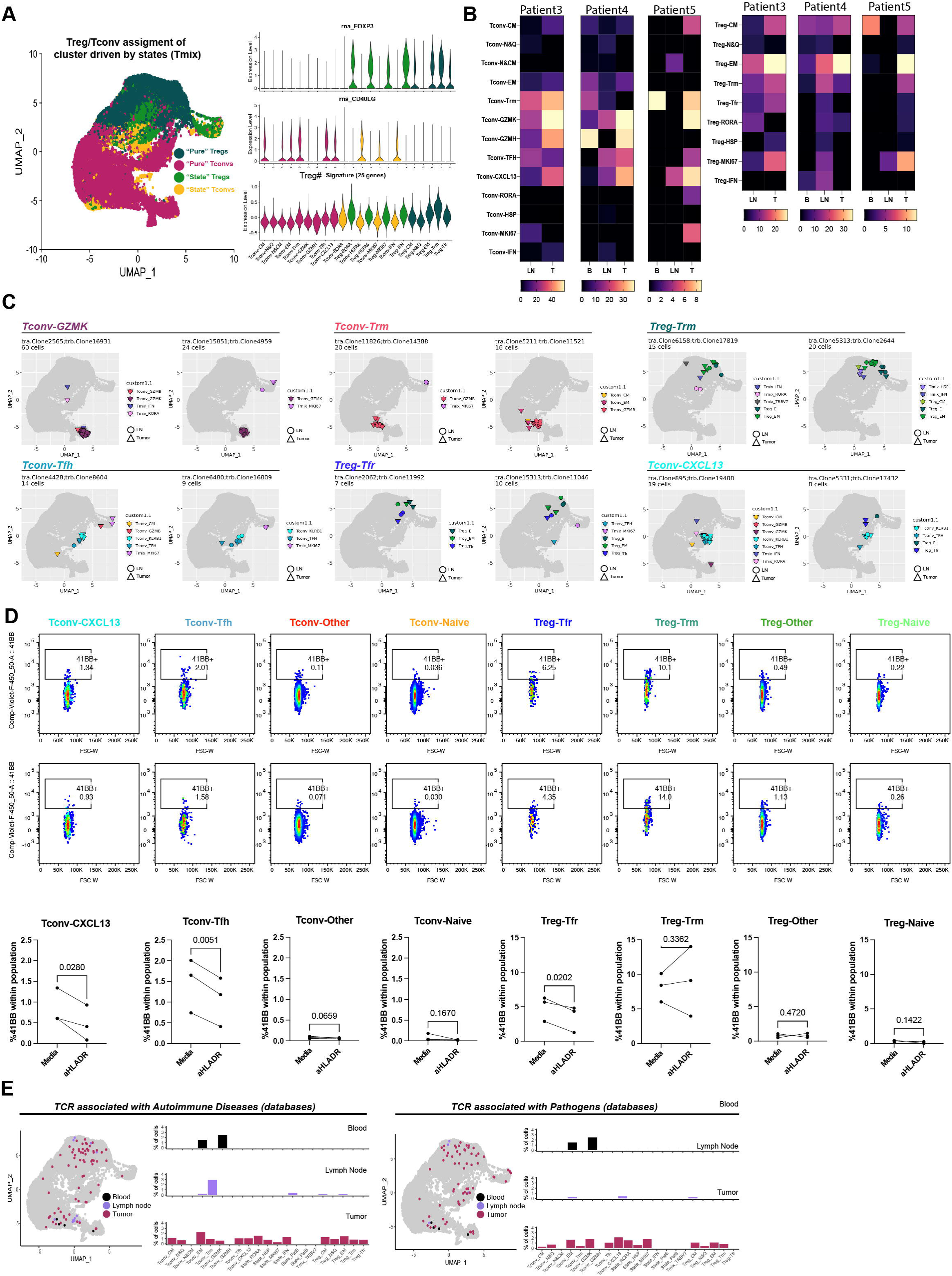
TCR analysis of CD4⁺ T cell populations across tissues. **(A)** Reassignment of state-driven cells (States) into Treg or Tconv clusters. **Left:** UMAP projection of transcriptomic data, colored by cluster group: pure Treg, pure Tconv, state-driven Treg (Treg mix), and state-driven Tconv (Tconv mix). **Right:** Violin plots displaying FOXP3 and CD40LG gene expression, along with Treg signature scores. **(B)** Heatmaps of the Gini Index per cluster/tissue across individual patients. **(C)** UMAP visualization of the scRNA-seq dataset illustrating the distribution of cells by individual expanded clones across tissues. Shapes represent tissue origin, and colors indicate cluster classification. **(D)** In vitro antigen presentation assay: Total metastatic LN cell suspensions were cultured ± anti-HLA-DR blocking antibody (10 µg/mL) for 16 hours. Flow cytometry was used to assess subpopulations and quantify 4-1BB expression, a marker of recent antigen recognition. **Top:** Representative dot plots of 4-1BB expression in the indicated subpopulations, with percentages shown. **Bottom:** Scatter plot showing corresponding quantifications. Statistical significance was determined using a paired t-test (n = 4 NSCLC patient lymph nodes; p-values indicated). **(E) Left:** UMAP projections of cells identified in external databases (VDJdb et McPAS), colored by tissue. **Right:** Bar plots showing the percentage of these cells across clusters by tissue.

**Supplementary Fig. S4:**
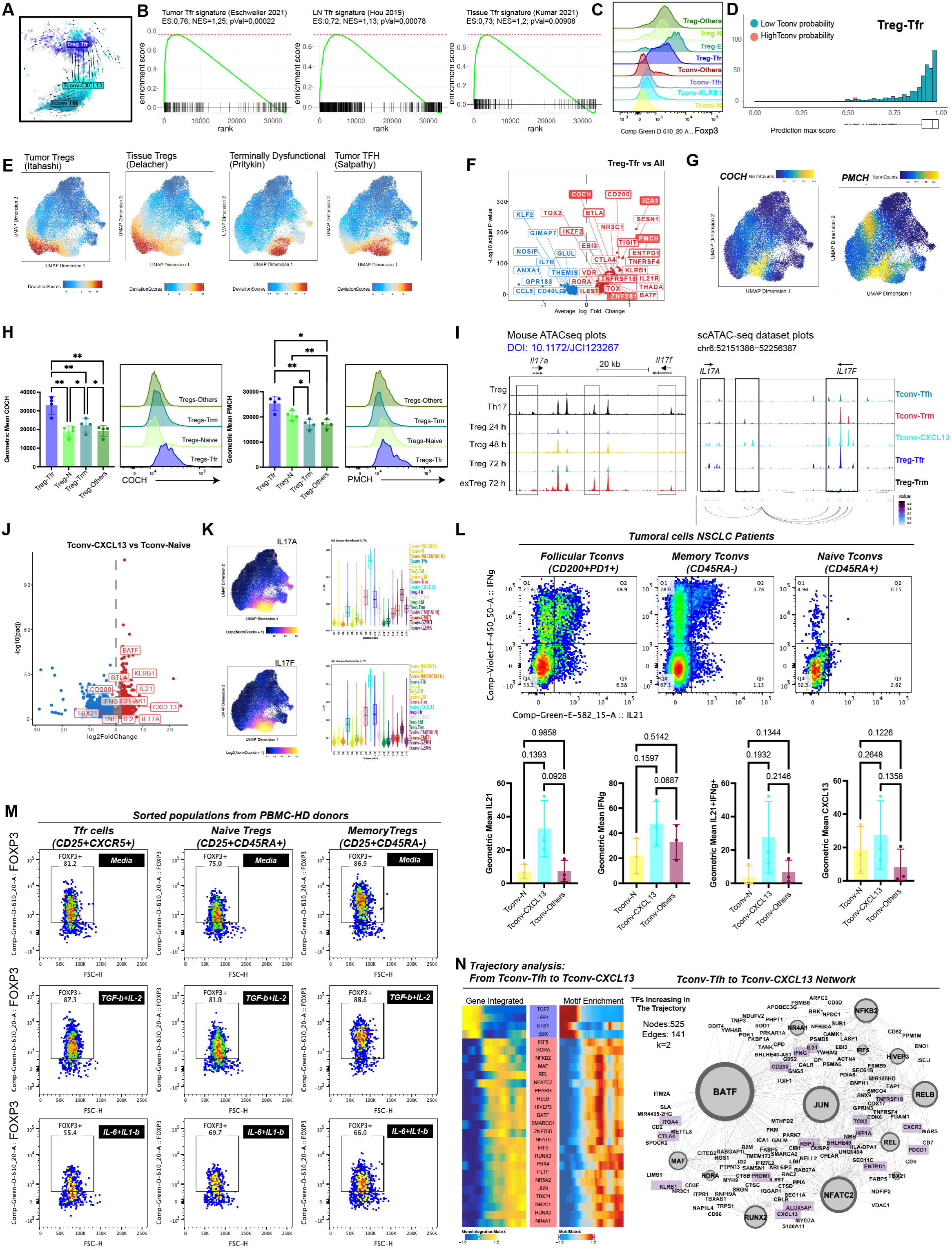
Integrated analysis reveals cell subsets transitions. **(A-D)** scRNA-seq analysis **(A)** scRNA-seq Velocity analysis: UMAP plot showing cells from the scRNA-seq dataset with RNA velocity vectors in lymph node and tumor cells. Streamlines depict directional flow for Tconv-Tfh, Tconv-CXCL13, and Treg-Tfr subsets. (**B)** Gene set enrichment analysis (GSEA) plot showing the Normalized Enrichment Score (NES) and p-values of Treg-Tfr cluster for publicly available signatures of Tfr cells. **(C)** Representative histograms showing FACS FOXP3 staining of sorted subpopulations. **(D)** Treg and Tconv label reassignment of Treg-Tfr cells: Histogram illustrating the number of cells by prediction score, classified as Tregs (green) or Tconvs (red) using the transfer label function within the Treg-Tfr cluster. **(E-G)** scATAC-seq analysis. **(E)** UMAP projection of per-cell Peak-signature enrichment public signatures for tumor-Tregs, tissue-Treg cells, terminally dysfunctional cells, and tumor Tfh cells **(F-G)** ScATAC-seq differential expression analysis of Treg-Tfr cells compared to all other clusters. Volcano plots **(F)** and representative UMAP plots (**G**) highlighting differentially expressed genes (Log2FC > 0.2, adj.pVal < 0.05), including *COCH* and *PMCH*. **(H)** Representative FACS histograms and bar plot quantifications of the geometric mean of COCH (**left**) and PMHC (**right**) across CD4+ T cell Treg subpopulations. **(I)** Chromatin accessibility of IL-17A/F loci: Genome tracks depicting normalized peak accessibility at *IL17A-F* gene loci in a publicly available mouse dataset (**left**) and human tissue-imprinted clusters from this study (**right**). Peak-to-gene links (P2G) from integrated scRNA-seq data are visualized (bottom arcs), with correlation strength indicated by color. **(J)** Bulk RNA-seq analysis: Volcano plot showing differential gene expression between Tconv-CXCL13 and Tconv-N&Q subsets. Cytokines characteristic of Th17-like cells and other Tconv-CXCL13-specific genes are highlighted (Log2FC > 1.5, adj.pVal < 0.05). **(K)** scATAC-seq gene scores: UMAPs and violin plots displaying gene score levels for *IL17A* (**top**) and *IL17F* (**bottom**), genes typically expressed by Th17 cells. **(L)** FACS analysis: (**upper panel**) Representative dot plots showing IL-21 and IFN-γ expression within gated populations from NSCLC tumor suspensions upon 5h stimulation with PMA/Ionomycin with Golgi-stop/GolgiPlug. (**bottom panel**) Bar plots quantifying the percentage of cells expressing IL-21, IFN-γ, both cytokines, and CXCL13 across different populations (*n*=3). Statistical significance was determined by paired one-way ANOVA with Tukey’s multiple comparisons test (p-values indicated). **(M)** In vitro polarization assay: Treg-Tfr, Treg-N, and Treg-Trm were FACS-sorted from healthy donor PBMCs and stimulated 3 days with anti-CD3/CD28 under different cytokine conditions as indicated. Representative dot plots show the percentage of FOXP3+ cells within live cells. Statistical test: Two-way ANOVA (paired) with Tukey’s multiple comparisons post hoc test (* = p ≤ 0.0332). **(N)** Transition from Tconv-Tfh to Tconv-CXCL13. **Right panels** display heatmaps of correlated gene integration (GI) and transcription factor (TF) motif enrichment along the trajectory. **Left panels** displays TF-target gene network: Regulatory networks driving Tconv-Tfh differentiation to Tconv-CXCL13. Node size represents out-degree, with selected target genes highlighted in squares.

**Supplementary Fig. S5:**
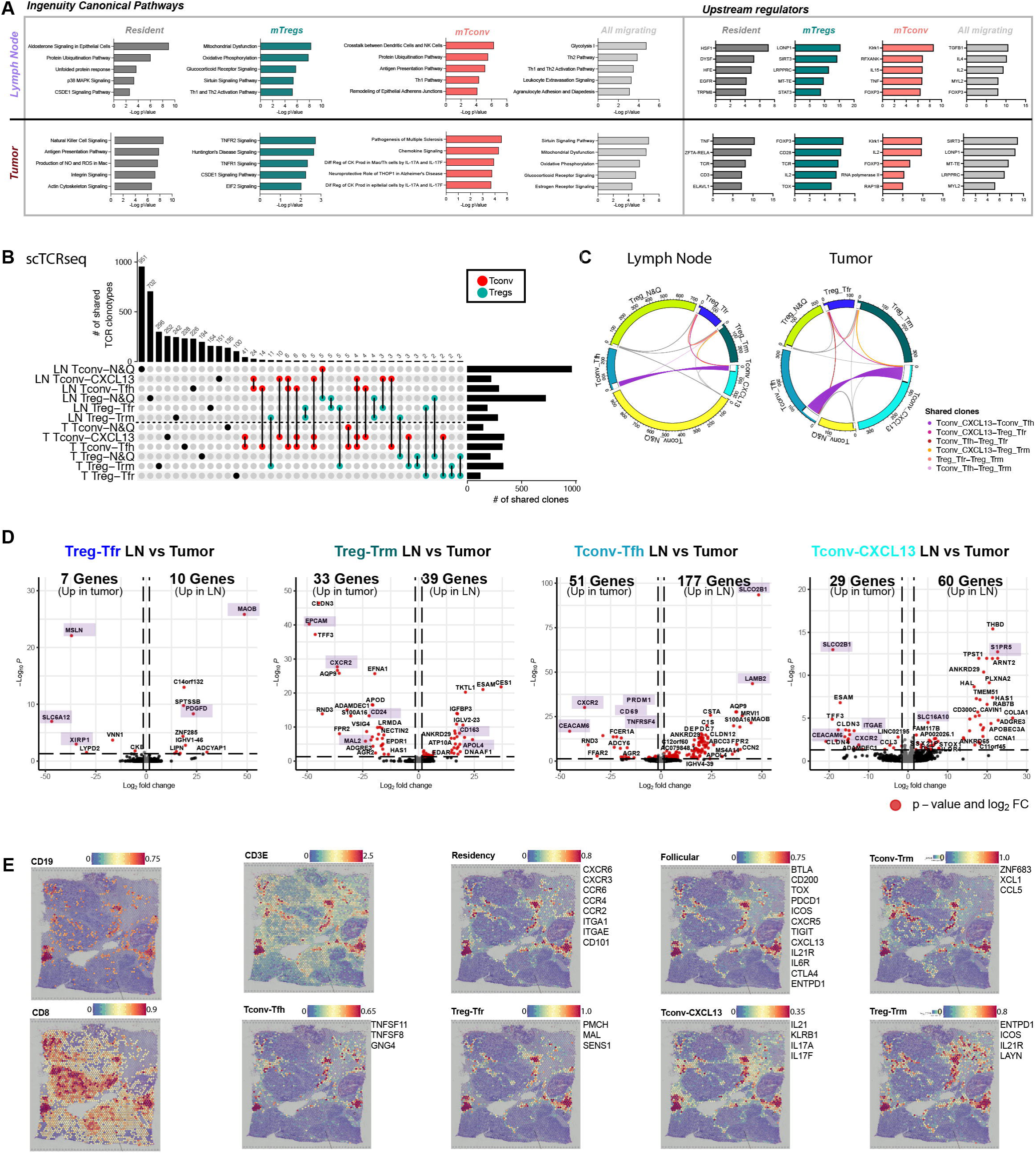
Integrated analysis reveals migration and tissue adaptation. **A-B)** Ingenuity Pathways Analysis (IPA): Canonical Pathways (**left**) and upstream regulators (**right**) analysis for the per tissue Log_2_ FC/FC plots comparing migrating and resident cells (Log2FC>0,2 and adj-pVal<0,05). **(B-C)** scTCR-seq analysis. TCR repertoire overlap across Treg-Trm, follicular and naïve Tregs and Tconv subpopulations split by tissue. (**B)** Upset plot display the number of clones/cells at each intersection. Lines (black) and circles (red for Tconvs and green for Tregs) indicate the populations involved in each comparison. Dot-line helps to visualize the separation between LNs and tumors. **(C)** Circos plots show clonotype distribution across populations in each tissue. Clonotypes shared between two subsets are highlighted in white, while those shared across multiple subsets appear in black. Key sharing patterns are color-coded. **(D)** Bulk RNA-seq analysis: Volcano plot showing differential gene expression between LN and Tumors from indicated subsets. Genes associated with migration, residency, and metabolisms are highlighted (Log2FC > 1.5, adj.pVal < 0.05). **(E)** Spatial transcriptomics analysis of human breast cancer tissue, showing the spatial distribution of CD3E mean expression and gene signature scores for resident and follicular general subsets, as well as specific Tconv-Trm, Tconv-Tfh, Treg-Tfr, Tconv-CXCL13, and Treg-Trm subset signatures using 10X Genomics Visium (public dataset, see material and methods). The genes used for each signature are listed on the right.

